# Estrogen receptor inhibition enhances cold-induced adipocyte beiging and glucose sensitivity

**DOI:** 10.1101/229591

**Authors:** Kfir Lapid, Ajin Lim, Eric D. Berglund, Jonathan M. Graff

## Abstract

Low estrogen states, exemplified by postmenopausal women, are associated with increased adiposity and metabolic dysfunction. We recently reported a paradox, in which a conditional estrogen receptor-alpha (ERα) mutant mouse shows a hyper-metabolic phenotype with enhanced brown/beige cell formation (“browning/beiging”). These observations led us to consider that although systemic deficiency of estrogen or ERα in mice results in obesity and glucose intolerance at room-temperature, cold-exposure might induce enhanced browning/beiging and improve glucose metabolism. Remarkably, studying cold-exposure in mouse models of inhibited estrogen signaling - ERαKO mice, ovariectomy, and treatment with the ERα antagonist Fulvestrant - supported this notion. ERα/estrogen deficient mice demonstrated enhanced cold-induced beiging, reduced adiposity and increased glucose sensitivity. Fulvestrant was also effective in diet-induced obesity settings. Mechanistically, ERα inhibition sensitized cell-autonomous beige cell differentiation and stimulation, including β3-adrenoreceptor-dependent adipocyte beiging. Taken together, our findings highlight a therapeutic potential for obese/diabetic postmenopausal patients.

## Introduction

Obesity and diabetes (“Diabesity”) afflict many in western civilizations, and are also a feature of estrogen deficiency^1,2.^ Hence, postmenopausal women are prone to show signs of increased adiposity and metabolic dysfunction^1,2^. Accordingly, obesity and metabolic dysfunction develop in pre-clinical estrogen-deficient models, including ERαKO and ovariectomized (OVX) female mice^3,4^. Hormone replacement therapy (e.g. estradiol administration) is successful in reversing metabolic dysfunction in rodent models and postmenopausal women^1,5^. Yet, hormone replacement therapy increases the risk of cardiovascular diseases and breast cancer^6,7^. Nonetheless, we recently uncovered an unanticipated function of ERα in systemic metabolism^8^. We found that mutant mice, with low estrogen signaling in the adipose lineage, display a lean phenotype, an improved glucose sensitivity and increased metabolic rates^8^. This paradoxical phenotype appeared secondary to an augmented brown/beige adipocyte differentiation, which appears to occur at the expense of white adipocyte differentiation^8^. Whereas white adipocytes primary function is to store excess energy intake, brown/beige adipocytes convert extra calories into thermal energy, thereby ameliorating diabesity^9-13^. Cold-exposure and sympathetic adrenergic signals are well-established stimuli that activate brown adipose tissue (BAT) and trigger formation of beige adipocyte^9-11^; the latter form within white adipose tissue (WAT). Indeed, cold temperatures are capable of reducing diabesity in rodents and human^9-11,14-16^. Since our aforementioned findings suggest that ERα regulates beiging/browning, we tested whether a browning/beiging stimuli, for example cold-exposure, may improve metabolic outcomes in mouse models of estrogen or estrogen receptor deficiencies. That is, estrogen receptor inhibition renders mice more susceptible to cold-induced browning/beiging effects, which do not normally occur at room temperature. To test our notion, we undertook a series of studies on independent and complementary models: whole-body ERαKO female mice, OVX females, and treatment with the potent estrogen receptor antagonist Fulvestrant (“Faslodex”™)^17^. In all models, we observed decreased adiposity and improved metabolic outcomes, which were secondary to enhanced cold-induced beiging and BAT activation. We also probed the effects in a primary beige cell culture, and demonstrated a cell-autonomous beiging following estrogen receptor inhibition. Finally, estrogen receptor inhibition appeared to enhance beiging also via β3-adrenoreceptor upregulation and activation.

## Methods

### Mice

WT strains include a mixed C57BL/6-129/SV background (for in vitro studies and acute Fulvestrant administration in vivo), ICR(CD1) (for chronic Fulvestrant administration in mice on normal chow) and C57BL/6 (for all other in vivo experiments). ERαKO mice were generously provided by Dr. Deborah J. Clegg^18^. Experiments were performed on 4-6-month-old female mice as indicated. The mice were housed in a temperature-controlled environment using a 12:12 light/dark cycle, and chow and water were provided ad libitum. Bilateral ovariectomy and sham operations were done as instructed in the standard operating procedure. The mice were fed either normal chow (4% fat, Harlan-Teklad) or high-fat diet (58% fat with sucrose, Research Diets). For cold experiments, mice were placed in a 6°C cold-chamber for 7 days or maintained at RT (23°C). Body temperatures were measured using a rectal probe. Fat/lean content/mass were measured using a minispec MQ10 NMR Analyzer (Bruker). Fulvestrant (40 mg/kg/injection, Sigma) or vehicle (DMSO) were dissolved in sunflower oil; acute administration – two intraperitoneal injections initiated 3 days prior to cold-exposure, or chronic administration - four intraperitoneal injections initiated 30 days prior to cold-exposure. CL-316,243 (1 mg/kg/day, Tocris Bioscience) was dissolved in H_2_O; three daily subcutaneous injections, mice were analyzed a day later. Mice were fasted for ~2 hr at RT prior to euthanization in most experiments, however, prior to glucose uptake assay and glucose/insulin tolerance tests, they were fasted for ~5 hr at RT. For glucose and insulin tolerance tests, 1.25 mg glucose (Sigma) or 0.3-0.75mU Humalog (Lilly)/g mouse weight were injected intraperitoneally; blood glucose levels were measured at the indicated intervals using Contour blood glucose monitoring (Bayer). Glucose uptake assay: jugular vein catheters were surgically implanted in mice a week prior to cold-exposure, using previously described procedures^19,20^. At the end of cold-exposure, mice were injected with a bolus of 13 μCi of [^14^C]-2-deoxy-D-glucose (2-DG) in the jugular vein catheter. Blood samples were obtained at intervals post-injection, after which mice were euthanized, and tissues were collected. Previously described procedures and calculations were used to determine plasma and tissue radioactivity, and to measure tissue-specific glucose uptake^18,19^. Other measurements were performed in the metabolic phenotyping core: sera insulin, triglycerides and cholesterol and liver triglycerides. All animal procedures were ethically approved by the UT Southwestern Medical Center institutional animal care and use committee.

### Cell culture

Stromal vascular (SV) fraction was obtained from subcutaneous adipose depots of two-month old mice as described^8,20,21^. Isolated SV cells were cultured in DMEM supplemented with 10% FBS, 100 units/ml penicillin, 100 mg/ml streptomycin and 25 ng/ml Amphotericin B (Sigma). White adipogenesis was induced as described^8,21^. Beige adipogenesis was induced similarly, except for an addition of 5 nM indomethacin (Sigma) and 2 nM T3 (Sigma)^20^. Fulvestrant (25 μM, Sigma), Estradiol (1 μM, Sigma), CL-316,243 (Tocris Bioscience), SR59230A (1 μM, Sigma), Propranolol (1 μM, Cayman Chemical) or vehicles (ethanol, H_2_O or DMSO in equivalent dilutions) were added to confluent cells with each media change. To induce thermogenic genes, cells were treated with 10 μM Forskolin (Sigma), 1 μM Norepinephrine (dissolved in H_2_O with 0.125 mM ascorbic acid, Sigma) or 1 μM CL-316,243 – 8 hr for RNA isolation or 24 hr for immunostaining. “Sub-optimal beige media” - IBMX and Dexamethasone (Sigma) were removed. Oil Red O staining, Nile Red staining and immunostaining were done as described^8,21^. Relative changes in cAMP levels were measured using cAMP-Glo™ Max Assay (Promega) following induction in PBS and IBMX (500 μM, Sigma) with or without 10 μM Forskolin for 15 min.

### Histology, immunohistochemistry and fluorescent immunostaining

Tissues were formalin-fixed, paraffin-embedded, sectioned and stained with haematoxylin and eosin (H&E) as described^21^. Immunohistochemistry and immunofluorescence were done as described^20,21^. Following de-paraffinization, the immunostaining procedure of paraffin-embedded tissue sections was similar to that of frozen tissues and cultured cells. Primary antibodies: rabbit anti-UCP1 (1:200, Abcam), rabbit anti-ERα (1:200, Abcam), rabbit anti-mouse β3-adrenergic-receptor (1:200, Abcam), mouse anti-PCNA (1:200, Millipore) and goat-anti-Perilipin (1:200, BD Biosciences). Secondary antibodies: Goat / Donkey anti-rabbit / mouse conjugated with AlexaFluor-488 / Cy3, and Donkey anti-goat conjugated with Cy5 (1:500, Jackson ImmunoResearch). DAPI (1:500, Fluka). Nile-red (1 μg/ml, Sigma). Bright-field and fluorescent images were collected on: Olympus inverted IX70 microscope, Olympus upright BX40 microscope, Leica inverted DMi8 microscope or Leica upright DM6 microscope.

### Quantitative teal-time PCR

Total RNA was extracted from adipose tissues or cultured cells as described^8,20^. cDNA synthesis and qPCR analysis of gene expression were done as described^8,20^. qPCR values were normalized to β-actin. Primer sequences are presented in Supplementary Table 1.

### Statistical analyses and data presentation

Statistical significance was assessed by two-tailed student’s t-test, area under curve and linear regression analyses. Error bars indicate S.E.M. Calculations were done and figures were generated using Microsoft Excel 2016, GraphPad Prism 7, ImageJ 1.5, CorelDraw X6 and ChemDraw 15.

## Results

### Whole-body ERαKO mice reduce adiposity and increase glucose sensitivity following cold-exposure

We previously reported that ERα plays alters adipose lineage specification: adipose lineage-restricted (stem cells to mature adipocytes) ERα mutants display increased beige adipocyte formation and BAT activity together with blunted white adipocyte formation^8^. These observations led us to consider that browning/beiging associated with ERα disruption in the adipose lineage might also be present in whole-body ERαKO mice under the right conditions. Induction of browning/beiging in ERαKO females, for example by cold-exposure^9^, is therefore predicted to result in fat loss and improved metabolism, despite their prominent obesity and metabolic dysfunction^3,22^. To test this possibility, we placed 4-5-month-old WT and ERαKO female mice in a cold-chamber for a week or kept them at room-temperature (RT). Cold-exposed ERαKO females significantly reduced body weight, fat content and fat mass, when compared to their counterparts at RT, whereas cold-exposed WT females did not (Fig. 1a-c). Further, the size and mass of subcutaneous and visceral fat depots were significantly decreased in cold-exposed ERαKO females (~3 fold-change) compared to cold-exposed WT females (Fig. 1d-f). Of note, cold-exposure did not appear to affect lean mass and weight of other organs (Fig. S1a-b). Since obesity is often associated with manifestation of diabetes, we tested whether the reduction in adiposity of ERαKO females is also associated with improvement of the metabolic profile. Although at RT, we detected hyperglycemia in ERαKO females, cold-exposure significantly lowered glucose levels, which were similar to the levels measured in cold-exposed WT females (Fig. 1g). We detected similar reductions in serum cholesterol and triglyceride levels in cold-exposed ERαKO females (Fig. 1h). We additionally performed glucose and insulin tolerance tests at RT or following a week of cold-exposure. Although at RT, ERαKO females demonstrated glucose intolerance and insulin resistance when compared to the WT counterparts, cold-exposure markedly increased their glucose and insulin sensitivities (Fig. 1i-j). Cold-exposure did not affect the glucose sensitivity of WT females, and only marginally their insulin sensitivity, however, WT females still performed better at these tests, compared to the ERαKO counterparts (Fig. 1i-j). Of note, high insulin levels remained intact following cold-exposure (Fig. S1c). Altogether, we found that upon cold-exposure, female ERαKO mice exhibited a significant reduction in adiposity and glucose levels together with increased glucose and insulin sensitivities. We had similar observations of reduction in adiposity and metabolic improvement in cold-exposed ERαKO males (not shown). These data suggest that in the absence of ERα mice are more sensitive to the metabolic-positive effects of cold, in contrast to their diabetic phenotype at RT.

**Figure 1.**
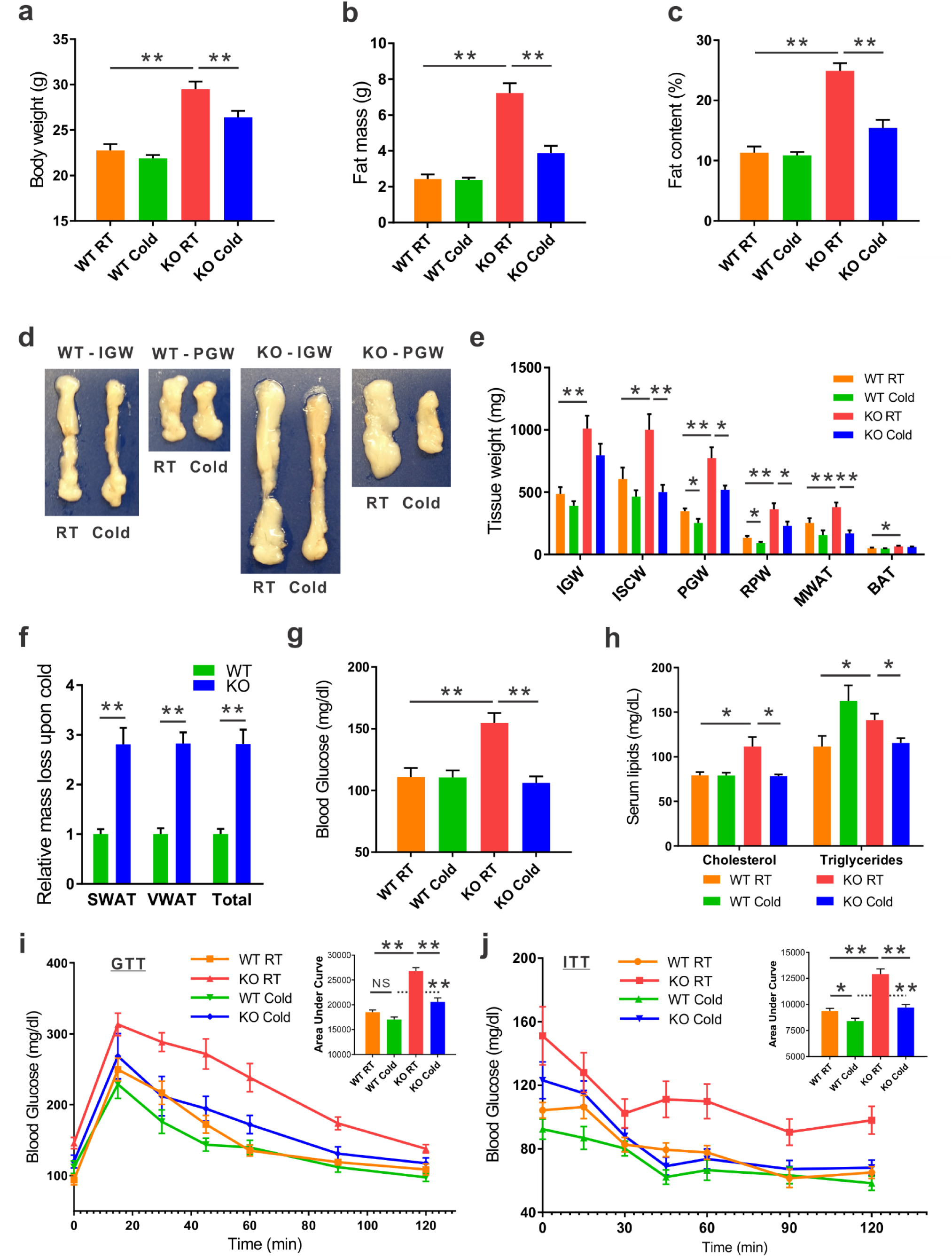
Whole-body ERαKO mice reduce adiposity and increase glucose sensitivity following cold-exposure. Four to five-month-old female ERαWT (WT) and ERαKO (KO) mice were subjected to cold-exposure (6°C) for 7 days or maintained at RT (23°C). **(a)** Body weight, n≥12. **(b)** Fat mass by NMR, n≥8. **(c)** Fat content by NMR, n≥8. **(d)** Representative photographs of inguinal (IGW) and perigonadal (PGW) white adipose depots of WT and KO females. **(e)** Weight of indicated fat depots in WT and KO females: SWAT - IGW and intrascapular (ISCW); VWAT - PGW, retroperitoneal (RPW) and mesenteric (MWAT); and intrascapular BAT, n≥8. **(f)** Relative fat mass loss in SWAT and VWAT compartments of WT and KO females were calculated according to white adipose depot weights in (e): SWAT = Cold(2*IGW+ISCW)-RT(2*IGW+ISCW); VWAT = Cold(2*PGW+2*RPW+MWAT)-RT(2*PGW+2*RPW+MWAT), n≥8. **(g)** Blood glucose levels of WT and KO females, n≥6. **(h)** Serum cholesterol and triglyceride levels of WT and KO females, n≥4. (**i-j**) Glucose tolerance (i) and insulin tolerance (j) tests were performed in WT and KO females at RT (a week prior to cold-exposure) and immediately after cold-exposure. Mice were fasted, i.p. injected with 1.25 g/kg glucose (i) or 0.3U/kg insulin (j), and their glucose levels were monitored, n≥8. Insets – areas under curve. Error bars indicate S.E.M. Statistical significance assessed by two-tailed student’s t-test, * p < 0.05, ** p <0.01.

### Whole-body ERαKO mice exhibit enhanced cold-induced beiging

According to our hypothesis, the cold-mediated effects on adiposity and glucose metabolism are secondary to an enhanced beiging in ERαKO females. At RT, obese ERαKO females were featured with large adipocytes in the subcutaneous WAT (SWAT) (Fig. 2a). At cold, histological examination of SWAT pointed to emergence of multilocular adipocytes, which were more abundant in ERαKO females compared to their WT counterparts (Fig. 2a). We did not notice any clear differences in the emergence of multilocular adipocytes in the visceral WAT (VWAT) at cold (Fig. 2b). At RT, interscapular classical BAT (or BAT) appeared “whiter” in ERαKO females, however, cold-exposure restored its appearance to a normal morphology (Fig. 2c). UCP1 expression is a hallmark of brown and beige multilocular adipocytes^9,13^. UCP1 immunohistochemistry of SWAT indicated an increased abundance of UCP1^+^ beige cells in cold-exposed ERαKO females as compared to cold-exposed WT females (Fig. 2d). A molecular evidence supported the suggested increased beiging: as compared to cold-exposed WT females, cold-exposed ERαKO females presented elevated mRNA levels of brown/beige cell markers in SWAT, including UCP1 (Fig. 2e). In contrast to SWAT, VWAT did not present an enhanced cold-induced beiging by gene expression (Fig. S2a). Nonetheless, we additionally detected an elevation of gene expression in BAT of ERαKO females (Fig. 2f). Even though we did not detect UCP1^+^ cells at RT (Fig. 2d), ERαKO females exhibited increased basal mRNA levels of UCP1 in SWAT (Fig. S2b, applies to other genes too – not shown), but neither in BAT nor in VWAT (not shown). Although increased BAT activity is often associated with a rise in body temperature^13^, cold-exposed ERαKO females reduced their body temperature in a similar manner as WT females (Fig. S2c). We had similar observations of enhanced cold-induced beiging in ERαKO males (not shown). Our data therefore suggest that reduced adiposity and improved glucose metabolism in ERαKO mice at cold are secondary to enhanced cold-induced browning/beiging.

**Figure 2.**
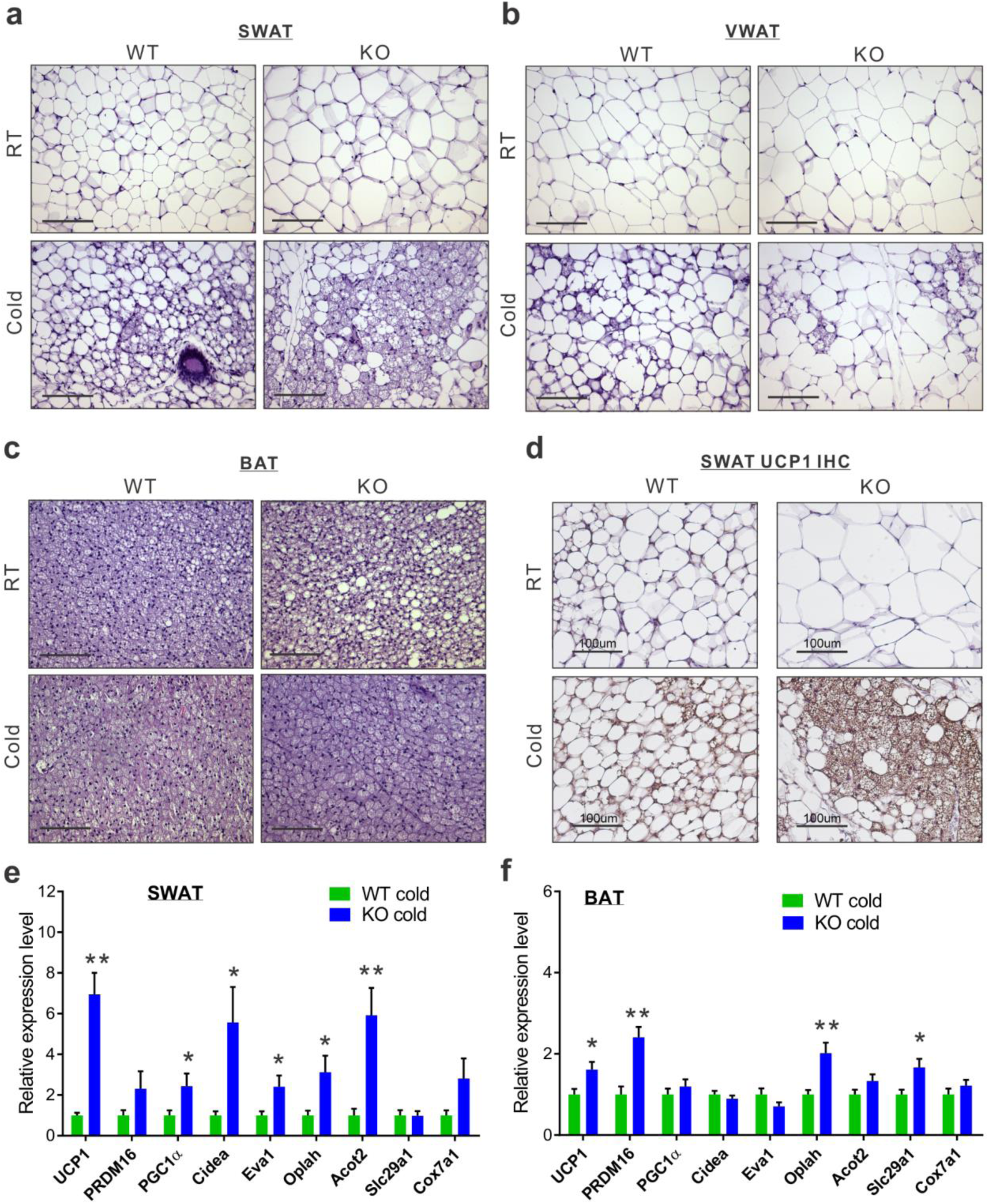
Whole-body ERαKO mice exhibit enhanced cold-induced beiging. Four to five-month-old female ERαWT (WT) and ERαKO (KO) mice were subjected to cold-exposure (6°C) for 7 days or maintained at RT (23°C). (**a-c**) Representative H&E-stained histological sections of SWAT (a), VWAT (b) and BAT (c).(**d**) UCP1 immunohistochemistry - representative sections of SWAT. (**e-f**) Relative mRNA levels, quantified by qPCR, of brown/beige adipocyte markers expressed in SWAT (IGW, e) and BAT (f) of WT and KO females at cold, n≥6. Scale bars = 100 μm. Error bars indicate S.E.M. Statistical significance assessed by two-tailed student’s t-test, * p < 0.05, ** p <0.01.

### Ovariectomized females reduce adiposity and increase glucose sensitivity following cold-exposure

In addition to a mouse model, lacking the receptor - ERα, a model lacking the ligand – estrogen, might serve as a complementary model to test the notion of enhanced cold-induced beiging. A common pre-clinical model in rodents for post-menopause is a surgical removal of the ovaries, consequently resulting in lower levels of circulating estradiol^23^. OVX female rodents gradually develop obesity and glucose intolerance, in consistence with metabolic manifestations of diabetic postmenopausal women^2,4,5,24^. Female mice were sham or OVX operated at 2-month of age. Three months later, allowing the OVX females to develop a diabetic phenotype, we placed sham and OVX mice in a cold-chamber for a week or kept them at RT. As expected, at RT, body weight, fat content and mass were higher in OVX females as compared to sham females (Fig. 3a-c). Following cold-exposure, OVX females significantly reduced body weight, fat content and fat mass, when compared to their counterparts at RT, while sham females only marginally (Fig. 3a-c). Further, the size and mass of subcutaneous and visceral fat depots were significantly decreased in cold-exposed OVX females (~4.5 fold-change) compared to cold-exposed sham females (Fig. 3d-f). Interestingly, the liver, which is fatty in OVX females^25^, reduced its weight following cold-exposure (Fig. 3g). Of note, cold-exposure did not appear to affect lean mass and other organs (Fig. S3a-b). Next, we evaluated whether the metabolic state of the OVX females could be improved in a similar manner to ERαKO females (Fig. 1). While at RT, we detected hyperglycemia in OVX females, cold-exposure significantly lowered glucose levels, which were similar to the levels measured in cold-exposed sham females (Fig. 3h). We detected similar reductions in serum cholesterol and triglyceride levels in cold-exposed OVX females (Fig. 3i). We additionally performed glucose and insulin tolerance tests at RT or following a week of cold-exposure. While at RT, OVX females demonstrated glucose intolerance when compared to the sham counterparts, cold-exposure markedly increased their glucose sensitivity (Fig. 3j). Unlike the positive effect of cold-exposure on the insulin resistance of ERαKO females (Fig. 1j), cold-exposure did not affect the insulin resistance of OVX females (Fig. S3c). Cold-exposure also did not affect the insulin sensitivity of sham females, and only marginally their glucose sensitivity, however, sham females still performed better at these tests, compared to the OVX counterparts (Fig. 3j and S3c). Of note, high insulin levels remained intact following cold-exposure (Fig. S3d). Altogether, we found that upon cold-exposure, OVX females exhibited a significant reduction in adiposity and glucose levels together with increased glucose sensitivity, suggesting that lower circulating estrogen sensitizes these mice to the metabolic-positive effects of cold.

**Figure 3.**
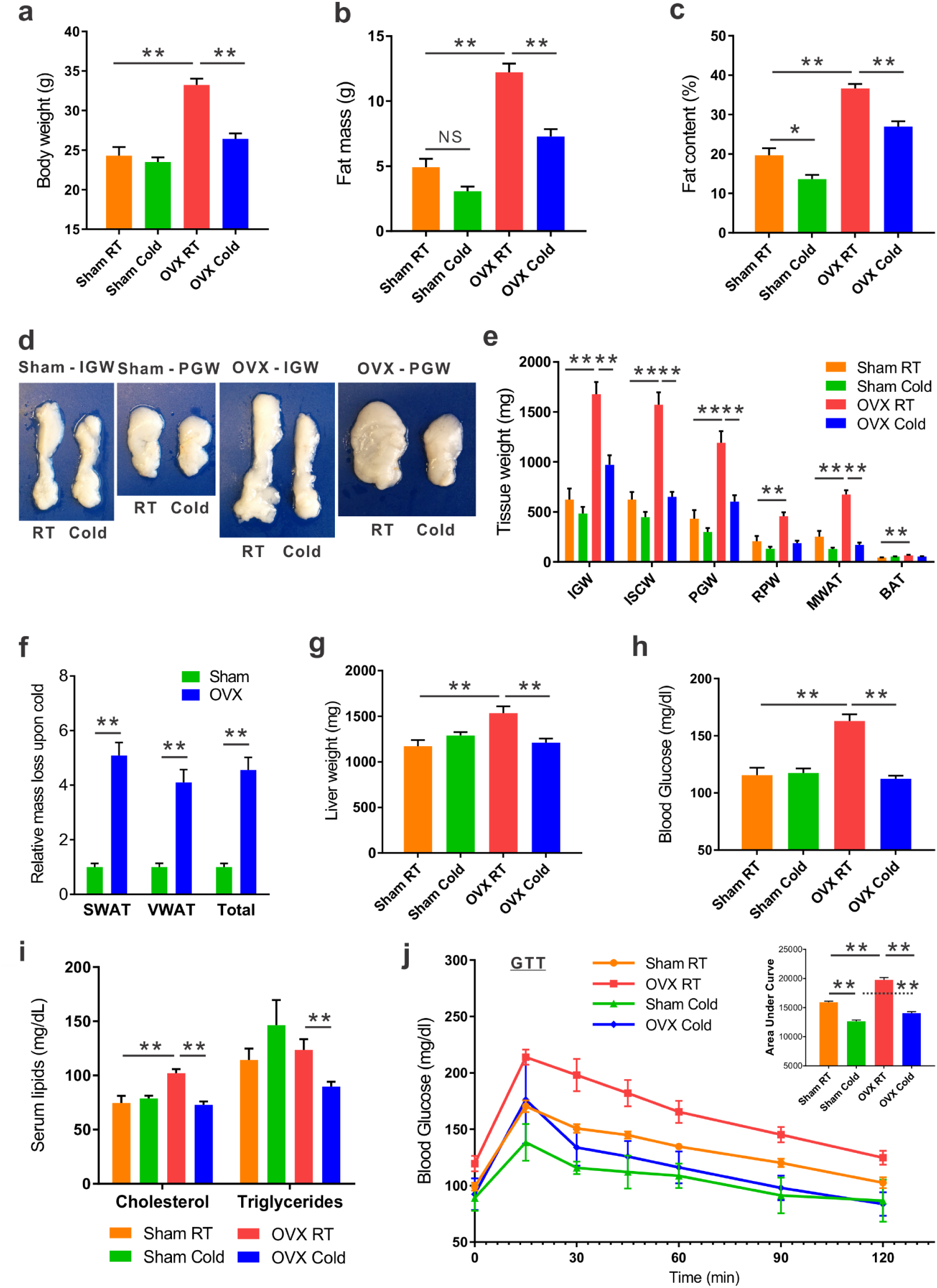
Ovariectomized females reduce adiposity and increase glucose sensitivity following cold-exposure. Two-month-old female mice underwent sham operation or ovariectomy (OVX), 3 months later both groups were either subjected to cold-exposure (6°C) for 7 days or maintained at RT (23°C). (**a**) Body weight, n≥11. (**b**) Fat mass by NMR, n≥6. (**c**) Fat content by NMR, n≥6. (**d**) Representative photographs of IGW and PGW adipose depots of Sham and OVX females. (**e**) Weight of indicated fat depots in Sham and OVX females: SWAT - IGW and ISCW; VWAT - PGW, RPW and MWAT; and intrascapular BAT, n≥8. **(f)** Relative fat mass loss in SWAT and VWAT compartments of Sham and OVX females were calculated according to white adipose depot weights in (e): SWAT = Cold(2*IGW+ISCW)-RT(2*IGW+ISCW); VWAT = Cold(2*PGW+2*RPW+MWAT)-RT(2*PGW+2*RPW+MWAT), n≥8. **(g)** Liver weights of Sham and OVX females, n≥8. (**h**) Blood glucose levels of Sham and OVX females, n≥8. **(i)** Serum cholesterol and triglyceride levels of Sham and OVX females, n≥8. **(j)** Glucose tolerance tests were performed in Sham and OVX females at RT (a week prior to cold-exposure) and immediately after cold-exposure. Mice were fasted, i.p. injected with 1.25 g/kg glucose, and their glucose levels were monitored, n≥9. Inset – areas under curve. Error bars indicate S.E.M. Statistical significance assessed by two-tailed student’s t-test, * p < 0.05, ** p <0.01, NS – not significant.

### Ovariectomized females exhibit enhanced cold-induced beiging

In line with findings in ERαKO females (Fig. 2), do we observe also enhanced cold-induced beiging in OVX females? At RT, obese OVX females were featured with large adipocytes in the SWAT and VWAT (Fig. 4a-b). At cold, histological examination of SWAT and VWAT pointed to emergence of multilocular adipocytes, which were more abundant in OVX females compared to their sham counterparts (Fig. 4a-b). At RT, BAT appeared “whiter” in OVX females, however, cold-exposure restored its appearance to a normal morphology (Fig. 4c). UCP1 immunohistochemistry of SWAT and VWAT indicated an increased abundance of UCP1^+^ beige cells in cold-exposed OVX females as compared to cold-exposed sham females (Fig. 4d and S4a). In line with enhanced cold-induced beiging in ERαKO females (Fig. 4e), cold-exposed OVX females presented elevated mRNA levels of brown/beige cell markers in SWAT as compared to cold-exposed sham females (Fig. 4e). We additionally detected an elevation of gene expression in VWAT and BAT of cold-exposed OVX females (Fig. S4b and 4f). As opposed to OVX females, enhanced cold-induced VWAT beiging was not evident in ERαKO females (Fig. 2b and S2a). Another difference between the two models is that OVX females showed similar basal mRNA levels of UCP1 in SWAT at RT (Fig. S4c). Despite increased browning/beiging at cold, OVX females reduced their body temperature in a similar manner as sham females (Fig. S4d). Aligned with reduction in liver weight (Fig. 3g), cold-exposure resulted in correction of estrogen deficiency-associated hepatosteatosis (Fig. 4g and S4e). Our data therefore suggest that reduced adiposity and improved glucose metabolism in OVX mice at cold are secondary to enhanced cold-induced browning/beiging.

**Figure 4.**
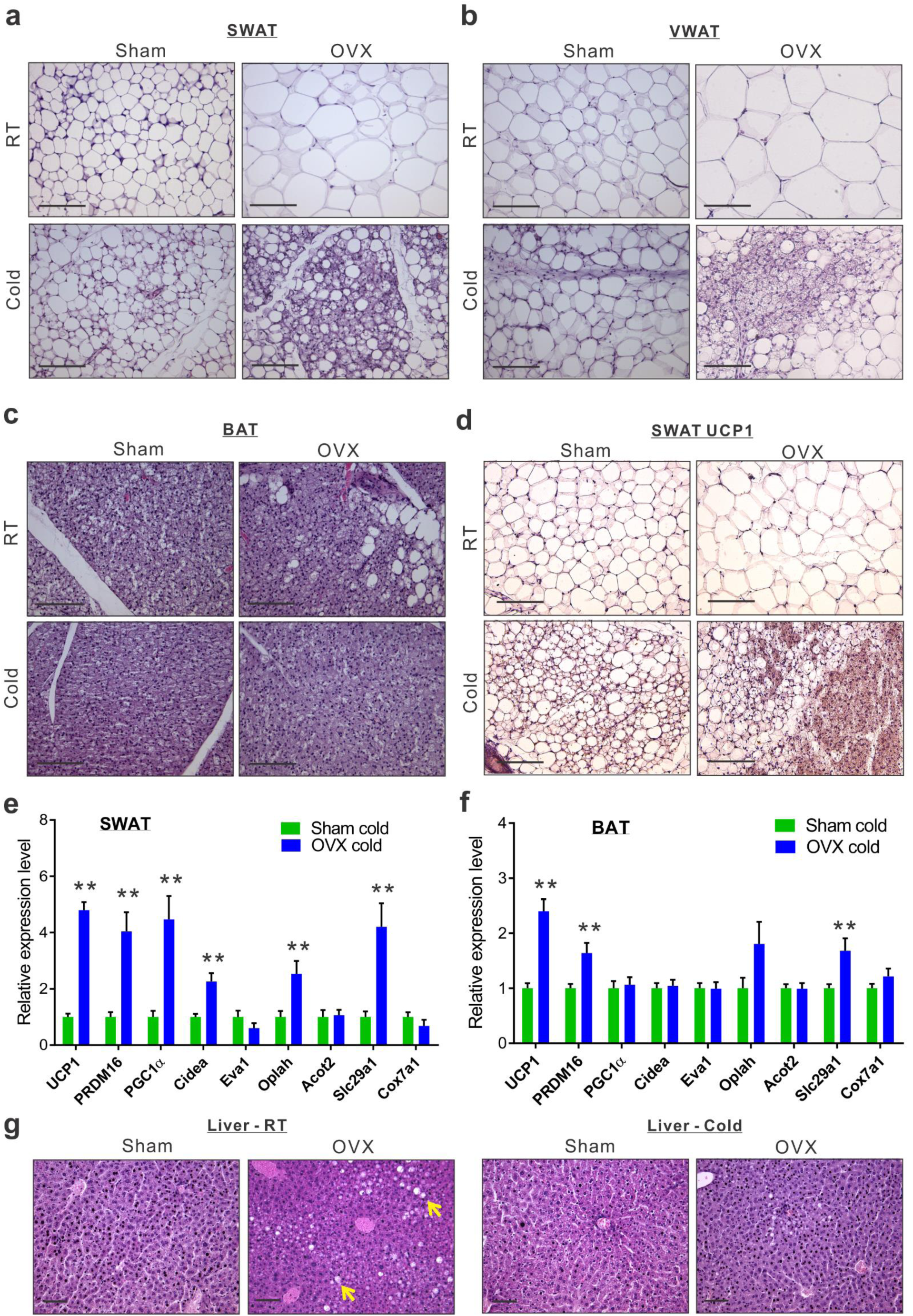
Ovariectomized females exhibit enhanced cold-induced beiging. Two-month-old female mice underwent sham operation or ovariectomy (OVX), 3 months later both groups were either subjected to cold-exposure (6°C) for 7 days or maintained at RT (23°C). (**a-c**) Representative H&E-stained histological sections of SWAT (a), VWAT (b) and BAT (c). (**d**) UCP1 immunohistochemistry - representative sections of SWAT. (**e-f**) Relative mRNA levels, quantified by qPCR, of brown/beige adipocyte markers expressed in SWAT (IGW) (e) and BAT (f) at cold, n≥6.(**g**) Representative H&E-stained histological sections of the liver. Arrows indicate fat deposits in OVX at RT. Scale bars = 100 μm. Error bars indicate S.E.M. Statistical significance assessed by two-tailed student’s t-test, * p < 0.05, ** p <0.01.

### Fulvestrant enhances beiging in vitro and in vivo

To test whether estrogen receptor inhibition enhances beiging in a cell-autonomous manner, we utilized a primary beige adipocyte culture. We isolated and cultured SWAT-derived stromal vascular (SV) cells, which are known to contain adipose progenitors^21^. Once the cells reached confluency, we exposed them to beige adipogenic media for a week and then activated them with Forskolin^20^. Since ERα-deficient progenitors have compromised adipogenesis^8^, we treated primary cells with the potent estrogen receptor antagonist Fulvestrant^17,26^. Unlike other selective estrogen receptor modulators (SERM), Fulvestrant is capable of ERα downregulation^27,28^, which we confirmed in vitro (Fig. 5a). Treatment with Fulvestrant during differentiation increased the number of UCP1^+^ beige cells in culture as compared to controls (Fig. 5b). Gene expression analysis of beige cell markers supported the notion of a cell-autonomous beiging (Fig. 5c). Notably, Fulvestrant did not affect adipogenesis in general (Fig. S5a), and was insufficient to trigger de-novo adipogenesis in non-adipogenic media (not shown). Fulvestrant was also incapable of converting white adipocytes to beige adipocytes (Fig. S5b), implying it enhances adipocyte beiging rather than inducing interconversion. Conversely, the ERα ligand, estradiol, has been reported to suppress in vitro white adipogenesis, and our results show that estradiol suppressed in vitro beiging well (Fig. 5b and S5c). Next, we tested whether estrogen receptor inhibition by Fulvestrant can also enhance cold-induced beiging in vivo. We acutely treated 4-month-old females with Fulvestrant or vehicle, and then placed them in a cold-chamber for a week or kept them at RT (Fig. 5d). Following cold-exposure, vehicle-treated and Fulvestrant-treated females did not show reduction in body weight (Fig. S5d). Nevertheless, reduction in adiposity was more significant in Fulvestrant-treated females as compared to vehicle-treated females (Fig. 5e-f and S5e). Acute Fulvestrant treatment did not appear to affect other organs (not shown). In accordance with cold effects on ERαKO females (Fig. 2a-b), acute Fulvestrant treatment augmented the emergence of multilocular adipocytes in SWAT at cold (Fig. 5g), but not in VWAT (Fig. S5f). At RT, acute Fulvestrant treatment had no effect on the histological morphology of WAT or BAT (Fig. 5f, S5f and not shown). UCP1 immunohistochemistry of SWAT at cold indicated an increased abundance of UCP1^+^ beige cells in Fulvestrant-treated females as compared to vehicle-treated females (Fig. 5h). Furthermore, at cold, Fulvestrant-treated females presented slightly elevated mRNA levels of brown/beige cell markers in SWAT and BAT as compared to vehicle-treated females (Fig. 5i-j). We had similar observations of enhanced cold-induced beiging in Fulvestrant-treated males (not shown). As a control experiment, acute administration of Fulvestrant in ERαKO mice did not result in any additive effect on cold-induced beiging (Fig. 5k and not shown). This supports the hypothesis that the effect of Fulvestrant on beiging is ERα-dependent. Taken together, ERα inhibition via Fulvestrant treatments mimics the effects of ERα absence in the enhancement of beiging in vitro and in vivo.

**Figure 5.**
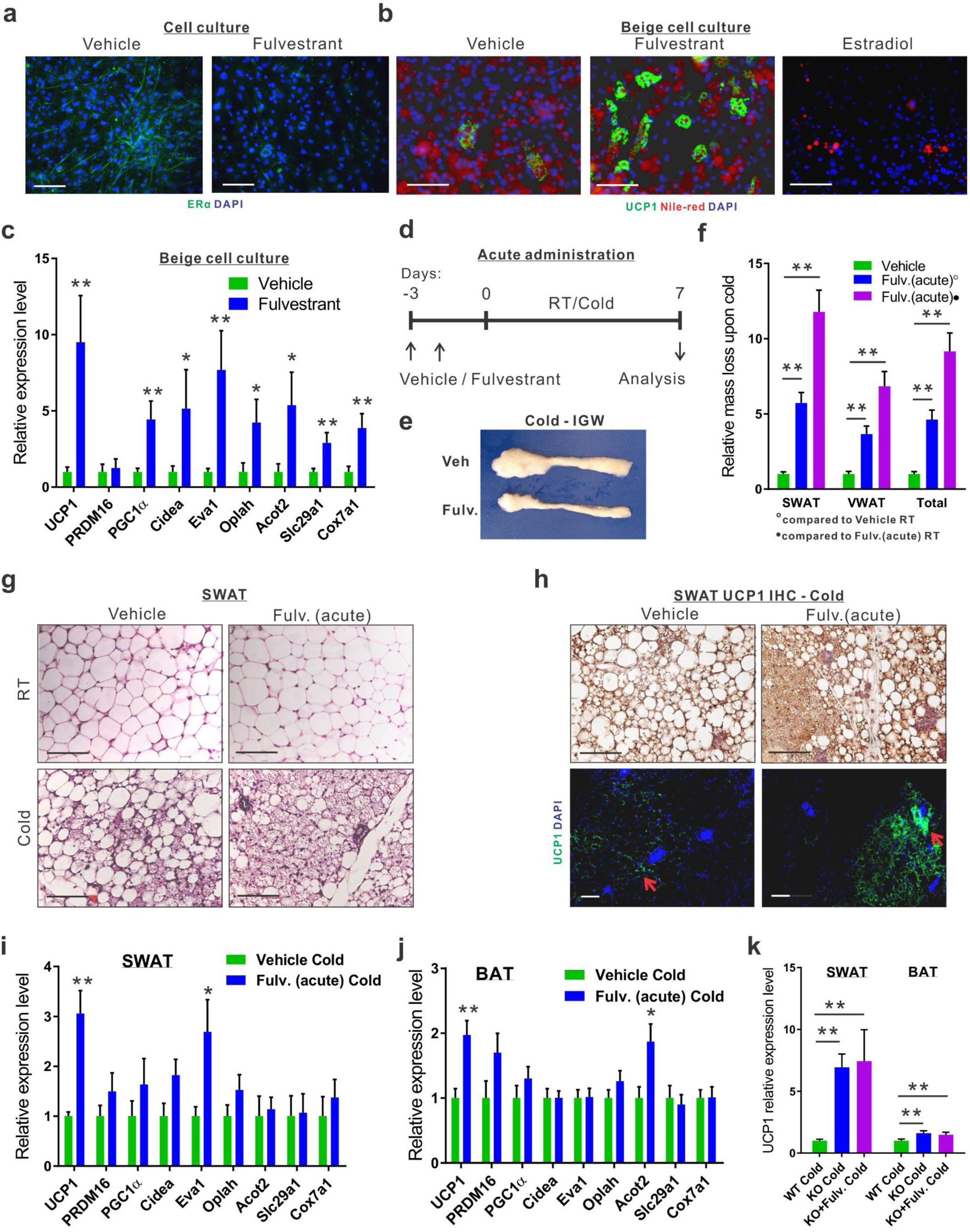
Fulvestrant enhances beiging in vitro and in vivo. (**a-c**) Stromal-vascular (SV) cells were isolated from SWAT of two-month-old females. (**a**) Confluent cells were treated daily with a vehicle or Fulvestrant for 48hr, and stained for ERα expression, using immunofluorescence. (**b-c**) Confluent cells were induced with beige adipogenic media in the presence of vehicles, Fulvestrant or Estradiol. A week later, beige cells were activated with Forskolin. Beiging was assessed by UCP1 immunostaining (b) or relative mRNA levels, quantified by qPCR, of brown/beige adipocyte markers, n≥7 (c). Nile Red stains lipid droplets. (**d-j**) Acute administration: Four-month-old WT females were given a vehicle or 40 mg/kg/injection Fulvestrant as described in (d), after they were subjected to cold-exposure (6°C) for 7 days or maintained at RT (23°C). (**e**) Representative photographs of IGW adipose depots of vehicle-treated and Fulvestrant-treated females at cold. (**f**) Relative fat mass loss in SWAT and VWAT compartments of vehicle-treated and Fulvestrant-treated females were calculated according to white adipose depot weights in (Fig. S5e): SWAT = Cold(2*IGW+ISCW)-RT(2*IGW+ISCW); VWAT = Cold(2*PGW+2*RPW+MWAT)-RT(2*PGW+2*RPW+MWAT), n≥7. **(g)** Representative H&E-stained histological sections of SWAT. (**h**) UCP1 immunohistochemistry (upper lane) and immunofluorescence (lower lane) - representative sections of SWAT at cold. Arrows indicate UCP1^002B^ cells. (**i-j**) Relative mRNA levels, quantified by qPCR, of brown/beige adipocyte markers expressed in SWAT (IGW) (i) and BAT (j) at cold, n≥5. (**k**) Vehicle or Fulvestrant administration in four to five-month-old ERαKO (KO) females in a similar manner to the WT females, as described in (d), after which they were subjected to cold-exposure (6°C) for 7 days. Relative mRNA levels, quantified by qPCR, of UCP1 in SWAT (IGW) and BAT, n≥5. Histological morphology confirms no difference in beiging between vehicle-treated and Fulvestrant-treated KO mice (not shown). Scale bars = 100 μm. Error bars indicate S.E.M. Statistical significance assessed by two-tailed student’s t-test, * p < 0.05, ** p <0.01.

### Chronic Fulvestrant treatment reduces adiposity and enhances cold-induced beiging

Acute Fulvestrant treatment moderately enhanced cold-induced beiging/browning in female mice. To test the beiging potential of a chronic Fulvestrant treatment, we used an outbred mouse strain of ICR(CD1) that is known to be heavier than the inbred C57Bl/6 and 129/SV strains used so far in this study^29^. The purpose is to demonstrate a generalized strain-independent beiging effect. We treated female mice with Fulvestrant or vehicle for a month (Fig. 6a), considering a long pharmacokinetic half-life of the drug^30^. Fulvestrant treatment was effective in downregulating ERα in WAT and BAT (Fig. S6a-c). We monitored the mice during the chronic Fulvestrant treatment - there were no notable changes in body weight, fat mass, fat content, lean mass (Fig. S6d-f), body temperature or food intake (not shown). Then, we placed them in a cold-chamber for a week or kept them at RT. Following cold-exposure, Fulvestrant-treated females significantly reduced body weight, fat content and fat mass, when compared to their counterparts at RT, while vehicle-treated females only marginally (Fig. 6b-d and S6d-f). Further, the size and mass of subcutaneous and visceral fat depots were significantly decreased upon cold-exposure in Fulvestrant-treated females (~2 fold-change) in comparison to vehicle-treated controls (Fig. 6e-g). Of note, Fulvestrant did not appear to affect lean mass and other organs (Fig. S6g-h). In contrast to the blood glucose lowering effect of cold-exposure in ERαKO and OVX females (Fig. 1g and 3h), we did not observe any differences in glucose levels (Fig. 6h); possibly because these mice are not hyperglycemic at RT. Although systemic glycemia did not change, WAT depots demonstrated increased glucose uptake capacity (2.3-3.7 fold-change) in Fulvestrant-treated females as compared to vehicle-treated females at cold conditions (Fig. 6i). Other organs showed no difference in glucose uptake capacity, including BAT (Fig. 6i). This might be due to the timing of the experiment or preference of fatty acid uptake by cold-activated brown adipocytes^16^. At cold, histological examination pointed at striking differences between Fulvestrant-treated females and vehicle-treated females – not only SWAT, but also VWAT displayed high abundance of multilocular adipocytes (Fig. 7a-b). At RT, chronic Fulvestrant treatment had no effect on the histological morphology of WAT or BAT (Fig. 7a-b and not shown). UCP1 immunohistochemistry of SWAT and VWAT at cold indicated an increased abundance of UCP1+ beige cells in Fulvestrant-treated females as compared to vehicle-treated females (Fig. 7c-d). Furthermore, at cold, Fulvestrant-treated females presented elevated mRNA levels of brown/beige cell markers in SWAT, VWAT and BAT as compared to vehicle-treated females (Fig. 7e-g). At RT, Fulvestrant-treated females exhibited increased basal mRNA levels of UCP1 in SWAT (Fig. S7a, applies to other genes too – not shown). Thus, chronic Fulvestrant pre-treatment enhanced cold-induced beiging to a larger extent than acute Fulvestrant pre-treatment. What’s more, we observed a physiological outcome that we did not observe in ERαKO and OVX females – while vehicle-treated females reduced their body temperature upon cold-exposure, Fulvestrant-treated females retained a higher body temperature (Fig. 7h). Monitoring the females at the course of cold-exposure showed that they were more resistant to the ambient cold temperatures (Fig. S7b), implying involvement of enhanced thermogenesis. In conclusion, chronic Fulvestrant treatment in female mice followed by cold-exposure leads to reduced adiposity, enhanced cold-induced beiging of WAT, increased glucose uptake in WAT and increased thermogenesis.

**Figure 6.**
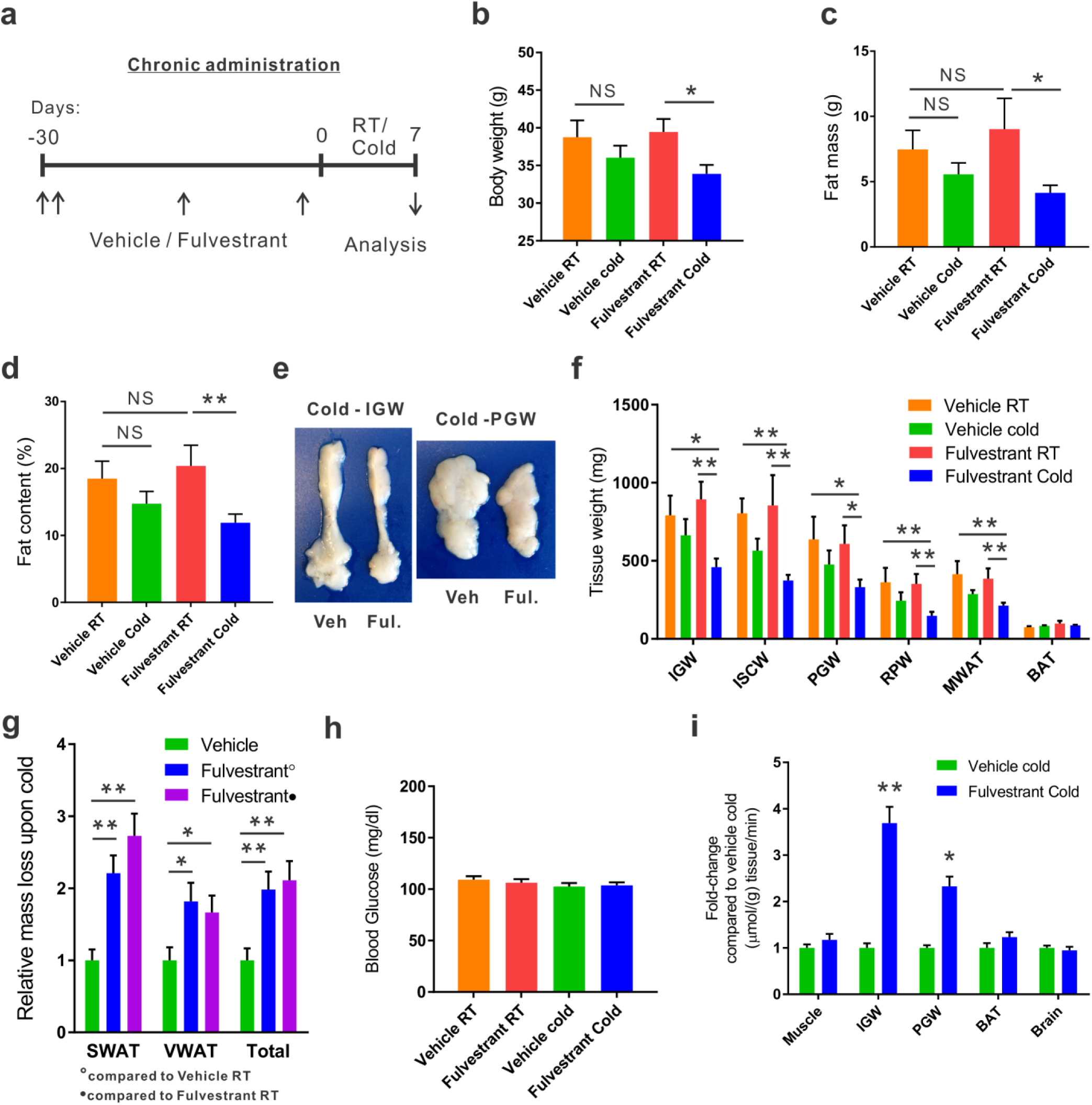
Chronic Fulvestrant treatment reduces adiposity. (**a-i)** Chronic administration: Four-month-old WT ICR(CD1) females were given a vehicle or 40 mg/kg/injection Fulvestrant for a month as described in (a), after they were subjected to cold-exposure (6°C) for 7 days or maintained at RT (23°C). (**b**) Body weight, n≥6. (**c**) Fat mass by NMR, n≥6. (**d**) Fat content by NMR, n≥6. (**e**) Representative photographs of IGW and PGW adipose depots of vehicle-treated and Fulvestrant-treated females at cold. (**f**) Weight of indicated fat depots in vehicle-treated and Fulvestrant-treated females: SWAT - IGW and ISCW; VWAT - PGW, RPW and MWAT; and intrascapular BAT, n≥6. (**g**) Relative fat mass loss in SWAT and VWAT compartments of vehicle-treated and Fulvestrant-treated females were calculated according to white adipose depot weights in (f): SWAT = Cold(2*IGW+ISCW)-RT(2*IGW+ISCW); VWAT = Cold(2*PGW+2*RPW+MWAT)-RT(2*PGW+2*RPW+MWAT), n≥6. (**g**) Blood glucose levels of vehicle-treated and Fulvestrant-treated females, n≥6. (**h**) In vivo glucose uptake assay, based on radiolabeled 2-DG administration, in vehicle-treated and Fulvestrant-treated females upon cold-exposure (see methods for further details), n≥5. Error bars indicate S.E.M. Statistical significance assessed by two-tailed student’s t-test, * p < 0.05, ** p <0.01, NS – not significant.

**Figure 7.**
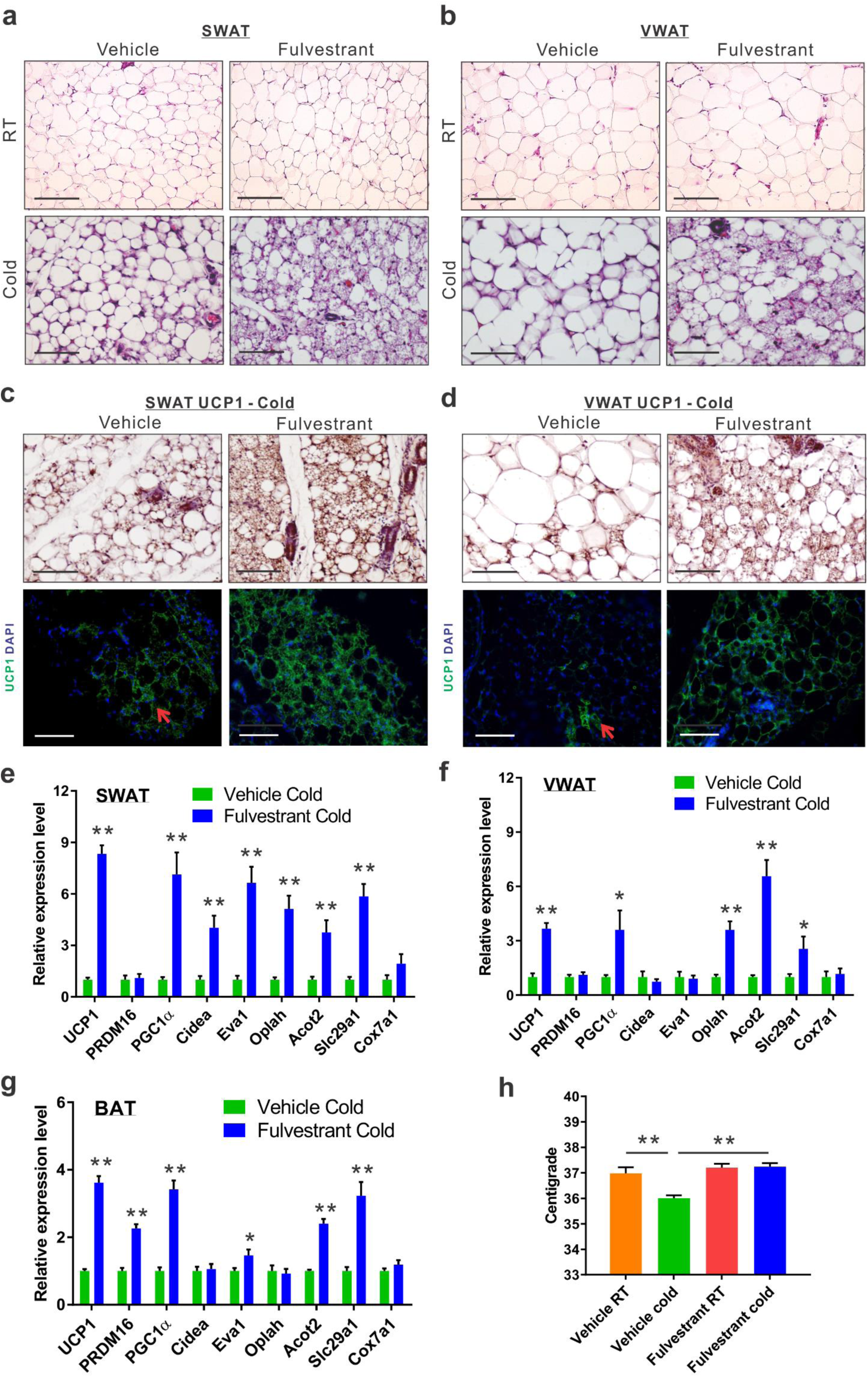
Chronic Fulvestrant treatment enhances cold-induced beiging. Chronic administration: Four-month-old WT ICR(CD1) females were given a vehicle or 40 mg/kg/injection Fulvestrant for a month as described in (Fig. 6a), after they were subjected to cold-exposure (6°C) for 7 days or maintained at RT (23°C). (a-b) Representative H&E-stained histological sections of SWAT (a) and VWAT (b). (c-d) UCP1 immunohistochemistry (upper lane) and immunofluorescence (lower lane) - representative sections of SWAT (c) and VWAT (d) at cold. Arrows indicate UCP1+ cells in vehicle-treated females. (e-g) Relative mRNA levels, quantified by qPCR, of brown/beige adipocyte markers expressed in SWAT (IGW) (e), VWAT (PGW) (f) and BAT (g) at cold, n≥8. (h) Body temperature (rectal probe) of vehicle-treated and Fulvestrant-treated females at RT and cold, n≥6. Scale bars = 100μm. Error bars indicate S.E.M. Statistical significance assessed by two-tailed student's t-test, * p < 0.05, ** p <0.01.

### Fulvestrant treatment improves the metabolism of diet-induced obese/diabetic mice

Our results using Fulvestrant propose that this pharmacological approach is potentially therapeutic. Whereas ERαKO and OVX females are obese and diabetic, the mice we used to test Fulvestrant so far were raised on normal chow (Fig. 5-7). Hence, we utilized a diet-induced diabesity mouse model fed with a high-fat high-sucrose diet (HFD). Aligning with the surgical OVX model, we fed 2-month-old females with HFD for 3 months, and during the last month, we administered Fulvestrant or a vehicle (Fig. 8a). As expected, HFD resulted in weight gain (Fig. S8a), and Fulvestrant administration had no effect on body weight, fat content, fat mass or lean mass when compared to vehicle treatment (Fig. 8b-c and S8a-c). Then, we placed them in a cold-chamber for a week or kept them at RT. Following cold-exposure, Fulvestrant-treated females reduced body weight and fat mass, when compared to vehicle-treated females at RT (Fig. 8b-c). The reduction in adiposity in females on HFD was moderate (Fig. 8c and S8b,d-e), unlike Fulvestrant treatment in females on normal chow (Fig. 6a-g). In accordance with observations in cold-exposed OVX females (Fig. 3g), Fulvestrant treatment in females on HFD reduced liver weight following cold-exposure (Fig. S8g), and the the treatment did not appear to affect other organs (not shown). Although Fulvestrant showed a moderate effect over adiposity, we observed a glucose-lowering effect of Fulvestrant in females on HFD, already evident at RT (Fig. 8d). In line with previous reports^15,16^, cold-exposure per se reduced the serum levels of glucose, cholesterol, triglycerides and insulin in mice on HFD (Fig. 8d and S8g-h), however, Fulvestrant treatment had no additive effect. We additionally performed glucose and insulin tolerance tests at RT or following a week of cold-exposure. Cold-exposure is known to increase glucose and insulin sensitivity in mice on HFD^15^, which we supported in evaluating vehicle-treated females (Fig. 8e-f). Nevertheless, Fulvestrant treatment in females on HFD had a significant synergistic effect in alleviating glucose intolerance at cold (Fig. 8e) and insulin resistance at RT (Fig. 8f), despite similar insulin levels (Fig. S8h). Next, we applied histology and gene expression analysis to females on HFD to evaluate browning/beiging. Histological examination and UCP1 immunoreactivity of SWAT at cold pointed to a scattered emergence of UCP1^+^ multilocular adipocytes, but they were not more abundant in Fulvestrant-treated mice (Fig. S8i and not shown). VWAT of females on HFD was resistant to cold-induced beiging, regardless of treatment (not shown). Already evident at RT, the “whiter” BAT, which characterized vehicle-treated females on HFD, was restored to a normal morphology in Fulvestrant-treated females on HFD (Fig. 8g). At cold, BAT morphology appeared even denser in Fulvestrant-treated females on HFD (Fig. 8g). On one hand, at cold, Fulvestrant-treated females presented only a moderate elevation in mRNA levels of brown/beige cell markers in SWAT as compared to vehicle-treated females (Fig. 8h). On the other hand, the elevation in the expression of these markers was more notable in BAT at cold (Fig. 8i). At RT, Fulvestrant-treated females exhibited increased basal mRNA levels of UCP1 in BAT, but not in SWAT (Fig. S8j, applies to other genes too – not shown). The histological and gene expression analyses suggest that HFD partially inhibited the enhancing effect of Fulvestrant on cold-induced WAT beiging, but not on BAT. Similarly to Fulvestrant-treated females on normal chow (Fig. 7h and S7b), Fulvestrant-treated females on HFD retained a higher body temperature (Fig. 8j and S8k). Moreover, in a similar manner to the observations in OVX females (Fig. 4g and S4e), we observed correction of HFD-associated hepatosteatosis in Fulvestrant-treated females following cold-exposure (Fig. 8k and S8l). In summary, our study demonstrates that estrogen receptor inhibition leads to reduced adiposity and improved glucose metabolism following cold-exposure, which are secondary to enhanced cold-induced beiging/browning.

**Figure 8.**
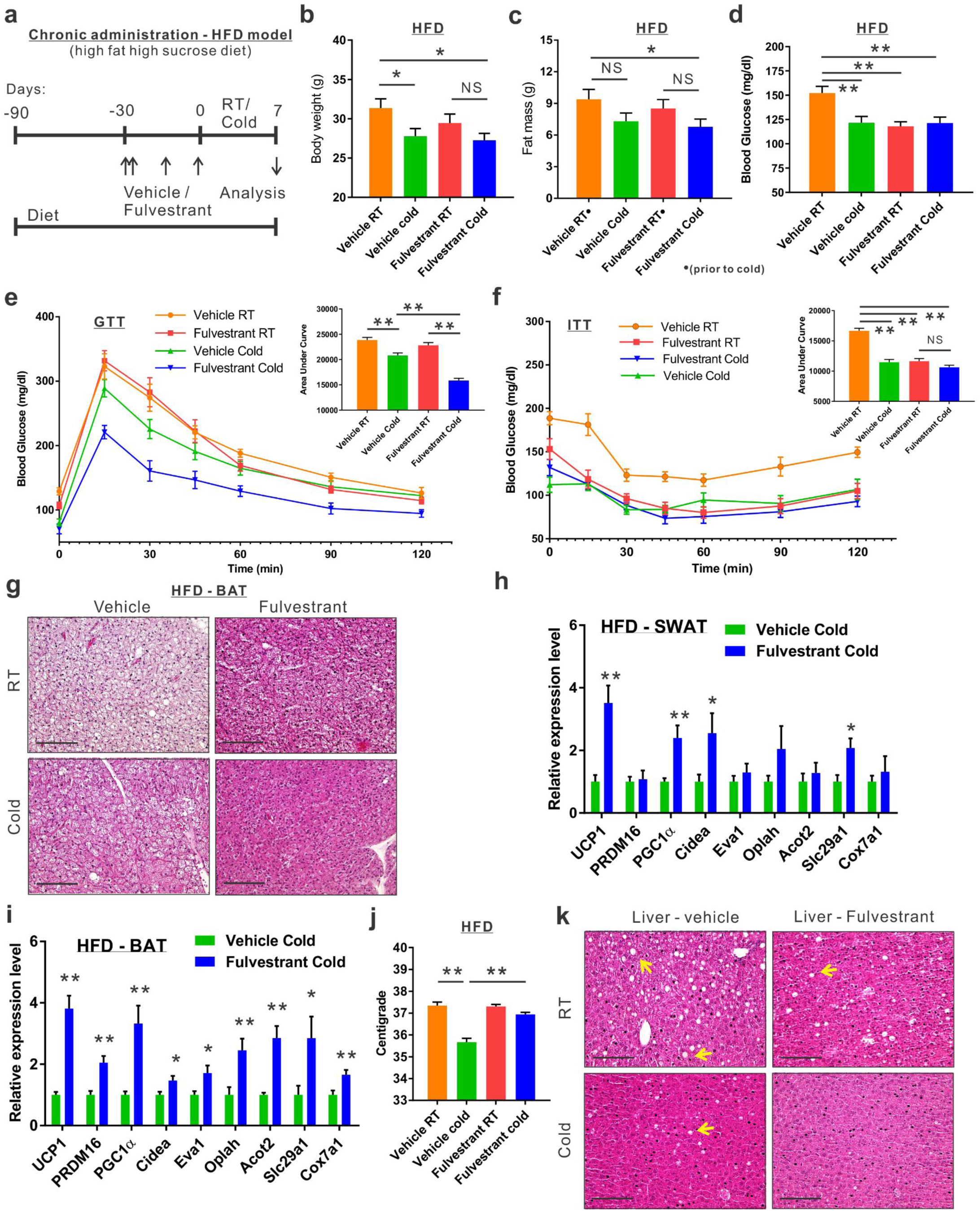
Fulvestrant treatment improves the metabolism of diet-induced obese/diabetic mice. (**a-k**) Chronic administration in a high-fat diet (HFD) model: At the age of two-month-old onwards, WT female mice were fed with high-fat high-sucrose diet. Two months later, the females were given a vehicle or 40 mg/kg/injection Fulvestrant for a month as described in (a), after they were subjected to cold-exposure (6°C) for 7 days or maintained at RT (23°C). (**b**) Body weight, n≥11. (**c**) Fat mass by NMR, n≥17. (**d**) Blood glucose levels of vehicle-treated and Fulvestrant-treated females on HFD, n≥7. (**e-f**) Glucose tolerance (e) and insulin tolerance (f) tests were performed in vehicle-treated and Fulvestrant-treated females on HFD at RT (a week prior to cold-exposure) and immediately after cold-exposure. Mice were fasted, i.p. injected with 1.25 g/kg glucose (e) or 0.75U/kg insulin (f), and their glucose levels were monitored, n≥8. Insets – areas under curve. (**g**) Representative H&E-stained histological sections of BAT. **(h-i)** Relative mRNA levels, quantified by qPCR, of brown/beige adipocyte markers expressed in SWAT (IGW) (h), and BAT (i) at cold, n≥6. (**j**) Body temperature (rectal probe) of vehicle-treated and Fulvestrant-treated females on HFD at RT and cold, n≥7. (**k**) Representative H&E-stained histological sections of the liver. Arrows indicate fat deposits in females on HFD, particularly vehicle-treated females at RT.Scale bars = 100 μm. Error bars indicate S.E.M. Statistical significance assessed by two-tailed student’s t-test, * p < 0.05, ** p <0.01.

### Estrogen receptor inhibition promotes beiging via β3-adrenoreceptor

Based on our observations, estrogen receptor inhibition enhanced cold-induced beiging. Adipocyte beiging is associated with multiple signaling pathways, predominantly cAMP-mediated pathways^31^. Indeed, Fulvestrant treatment in vitro elevated basal cAMP levels in differentiated beige cells and Forskolin-induced cAMP levels in undifferentiated cells (Fig. 9a and Fig. S9a), suggesting ERα inhibition “primes” cells to beiging. One of the key stimulants of cAMP-mediated signaling in adipocytes is the sympathetic nervous system, primarily via β-adrenoreceptors^9,13,31^. We therefore examined whether β-adrenoreceptors are upregulated in response to in vitro Fulvestrant treatment. We detected a subtype-restricted upregulation of β3-adrenoreceptor (AdRβ3) in beige cells, whether Fulvestrant is added during differentiation or after they differentiate (Fig. 9b-d). Fulvestrant treatment of non-induced undifferentiated cells also resulted in an upregulation of β-adrenoreceptors, but not in a subtype-restricted manner (Fig. S9b-d). We should emphasize that the mRNA levels of UCP1 and AdRβ3 in undifferentiated cells were inferior to that of differentiated beige cells (Fig. S9d), and that UCP1 was absent at the protein level in Fulvestrant-treated undifferentiated cells (not shown). PPARγ2 is an essential transcription factor that drives white/beige adipocyte differentiation^32-35^. In differentiated beige cells, Fulvestrant did not affect PPARγ2 expression (Fig. 9b-c), however, Fulvestrant increased its expression in undifferentiated cells (Fig. S9b). Together with no effect on cell proliferation (Fig. S9e) or adipocyte formation (Fig. S5a-b and not shown), these results suggest that Fulvestrant gives a “head-start” in the beiging process. Notably, it is insufficient to trigger beiging by itself, and requires external signals provided by the beige adipogenic media. Could Fulvestrant enhance beiging via AdRβ3 in a similar manner to cold-induced beiging? Since Fulvestrant pre-treatment upregulated AdRβ3 expression (Fig. 9d and S9c), we first pre-treated cells with Fulvestrant or a vehicle, and then stimulated the cells with the specific AdRβ3 agonist CL-316,243 in the absence of Fulvestrant. Indeed, Fulvestrant pre-treatment increased the number of UCP1^+^ beige cells in response to stimulation by CL-316,243, whether it is added during differentiation (Fig. 9e) or prior to induced differentiation (Fig. S9f). In other words, Fulvestrant pre-treatment enhanced CL-316,243-induced beiging in vitro. To further support an AdRβ3-mediated effect, we stimulated cultured beige cells by the natural ligand, Norepinephrine (NE). Not only that Fulvestrant enhanced NE-induced beiging, co-treating beige cells with the specific AdRβ3 antagonist SR59230A inhibited NE-induced beiging, regardless of treatment (Fig. 9f). Co-treating the cells with a pan-AdRβ blocker propranolol completely abolished NE-induced beiging (not shown). Next, we tested whether AdRβ3 stimulation in Fulvestrant-treated females and ERαKO females results in enhanced beiging in a similar manner to cold-exposure. Fulvestrant-treated WT females and ERαKO females showed an upregulated AdRβ3 expression in SWAT as compared to their control counterparts (Fig. S9g-h). We pre-treated 4-month-old female mice with vehicle or Fulvestrant, followed by administration of CL-316,243 injections (Fig. 9g). Following CL-316,243 administration, females reduced adiposity, however, reduction in adiposity was more significant in Fulvestrant-pre-treated females as compared to vehicle-pre-treated females (Fig. S9i). We then evaluated SWAT beiging by histological morphology, UCP1 immunostaining and gene expression analysis. Strikingly, Fulvestrant-pre-treated females demonstrated an enhanced beiging response to CL-316,243 administration as compared to vehicle-pre-treated females (Fig. 9h-i and S9j). Despite significant elevation in the expression of brown/beige cell markers in SWAT (Fig. S9j), Fulvestrant pre-treatment did not result in a profound effect on BAT (Fig. S9k). Finally, ERαKO females demonstrated an enhanced beiging effect in response to CL-316,243 administration (Fig. 9j-k and not shown). In summary, estrogen receptor inhibition enhances adipocyte beiging in vitro and in vivo via AdRβ3 upregulation and activation.

**Figure 9.**
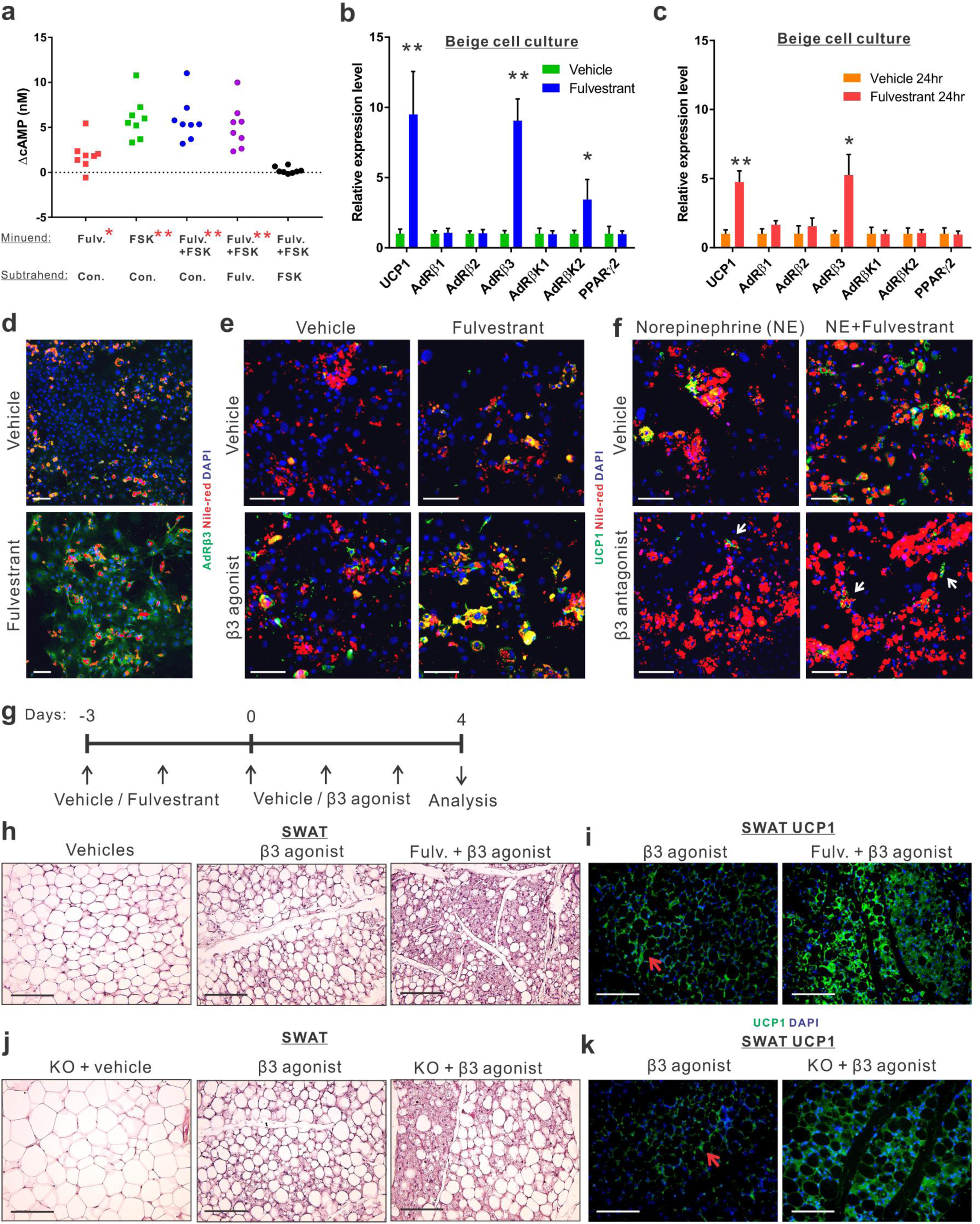
Estrogen receptor inhibition promotes beiging via β3-adrenoreceptor. (a-f) SV cells were isolated from SWAT of two-month-old WT females. Confluent cells were induced with beige adipogenic media in the presence of vehicle or Fulvestrant. A week later, the experiments below were performed. (**a**) Relative changes in cAMP levels upon induction with or without Forskolin. Every dot represents an average of technical triplicates of one biological sample. The relative changes in cAMP levels are calculated as differences between treatments, n≥8. **(b-c)** Relative mRNA levels, quantified by qPCR, of UCP1, β-adrenergic receptors, β-adrenergic receptor kinases and PPAR_γ_2. (**b**) Vehicle-treated or Fulvestrant-treated beige cells were activated with Forskolin for 8hr, n≥7. (**c**) Vehicle or Fulvestrant were not added during differentiation, but rather to mature beige cells for 24hr without Forskolin, n≥5. **(d-f)** Vehicle or Fulvestrant were added during differentiation. Nile Red stains lipid droplets. (**d**) Vehicle-treated or Fulvestrant-treated beige cells were immunostained for AdRβ3 expression. (**e**) Vehicle-treated or Fulvestrant-treated beige cells were activated or not with CL-316,243 (β3 agonist) for 24hr, and then immunostained for UCP1 expression. (**f**) Vehicle-treated or Fulvestrant-treated beige cells were activated with Norepinephrine (NE) for 24hr; concomitantly co-treated with a vehicle or SR59230A (β3 antagonist), and then immunostained for UCP1 expression. **(g-i)** Four-month-old WT females were pre-treated with a vehicle or 40 mg/kg/injection Fulvestrant, followed by administration of 1 mg/kg/day β3 agonist as described in (g). Representative histological sections of SWAT, which were either H&E-stained (h) or immunostained for UCP1 expression (i). (**j-k**) Four to five-month-old ERαWT or ERαKO (KO) females were treated with β3 agonist as described in (g). Representative histological sections of SWAT, which were either H&E-stained (j) or immunostained for UCP1 expression (k). Arrows indicate UCP1^+^ cells. Scale bars = 100 μm. Error bars indicate S.E.M. Statistical significance assessed by two-tailed student’s t-test, * p < 0.05, ** p <0.01. In (a), results are based on a matched standard curve and a linear regression analysis.

## Discussion

Mouse models of estrogen or ERα deficiencies often develop diabesity, characterized by high fat mass, glucose intolerance, insulin resistance, hyperlipidemia and hepatosteatosis^2-4,22,24,25^. In this study, we show that cold-exposure led to reduced adiposity and improved metabolic profile in ERαKO females, OVX females and Fulvestrant-treated females on HFD (summarized in Fig. 10a-b). Supported by our results, cold-exposure by itself is capable of improving glucose metabolism in obese/diabetic rodent models^15^. Yet, our results imply that under circumstances of estrogen receptor inhibition, mice experience a boosted improvement of glucose metabolism in response to cold-exposure. While at “normal” conditions of RT, ERαKO and OVX female mice were obese/diabetic, at cold, they were more susceptible to its metabolic-beneficial effects. In all these models, including regimens of Fulvestrant treatment in female mice on normal-chow, we detected an enhanced cold-induced beiging, predominantly in SWAT, according to histological morphology and gene-expression profiles. We noticed some differences in the manifestation of browning/beiging between these models. For example, cold-induced VWAT beiging occurred only in OVX females and following chronic Fulvestrant pre-treatment in mice on normal-chow. On the other hand, Fulvestrant pre-treatment in mice on HFD resulted in a mild cold-induced beiging, attaining the effects on BAT. While it is conceivable that the enhanced cold-induced beiging and BAT activity exert the observed beneficial effects on glucose metabolism, we cannot exclude that other tissues with ablated estrogen signaling play a cold-inducible role. Tissue-specific ERα mutants, such as deletions of ERα in mature adipocytes, hepatocytes, skeletal muscle and hypothalamic neurons, have been shown to partially develop characteristics of metabolic dysfunction^18,36-38^. Estrogen-dependent regulation of energy balance by the central nervous system is a key physiological contributor, and may explain in part the obese/diabetic phenotype of whole-body ERαKO and OVX mice^38-40^. Nonetheless, Fulvestrant, which does not appear to cross the blood-brain-barrier^26^, has an advantage over other SERMs in our understating of central vs. periphery roles of ERα. In support, cerebroventricular injections of Fulvestrant reduce systemic energy expenditure^41^, whereas intraperitoneal injections do not. What’s more, Fulvestrant downregulates ERα expression^27,28^, including in our study, and therefore serves as a reliable model for ERα deficiency. We presented a cell-autonomous beiging response under conditions of ERα deficiency by utilizing an isolated primary cell culture. We previously reported an enhanced beiging in mice with an adipose-lineage-specific deletion of ERα^8^, and herein we further validate this beiging effect in mice with systemic ERα-deficiency followed by stimuli of beiging/browning. These stimuli, elicited by cold-exposure or adrenergic signals, are necessary to amplify the downstream signaling triggered by ERα inhibition, which do not generate a beiging phenotype under non-stimulatory conditions (i.e. RT). This unknown downstream signaling is yet to be elucidated, however, our preliminary findings indicated upregulated cAMP levels, which are a driving force in beiging, as well as an increased AdRβ3 expression. Although AdRβ3 signaling triggers prominent beiging^13,31,42^, AdRβ3-KO mice and mice treated with AdRβ3 antagonist display normal cold-induced beiging^43,44^, suggesting AdRβ3 is redundant in the specific response to cold-exposure. The findings in this study are not contradictory, since they propose that following ERα inhibition, brown/beige adipocytes are more sensitive to activation – either by cold-exposure or by AdRβ3 agonist. This increased sensitivity may occur both at the immature and mature cell level (Fig. 10c), although this cellular mechanism should be further validated in vivo. Altogether, our results suggest that pre-clinical mouse models of post-menopause have a propensity to increased beiging and improved metabolism upon cold-exposure. Thus, we conclude that cold-exposure has a clinical potential in obese/diabetic postmenopausal patients.

**Figure 10.**
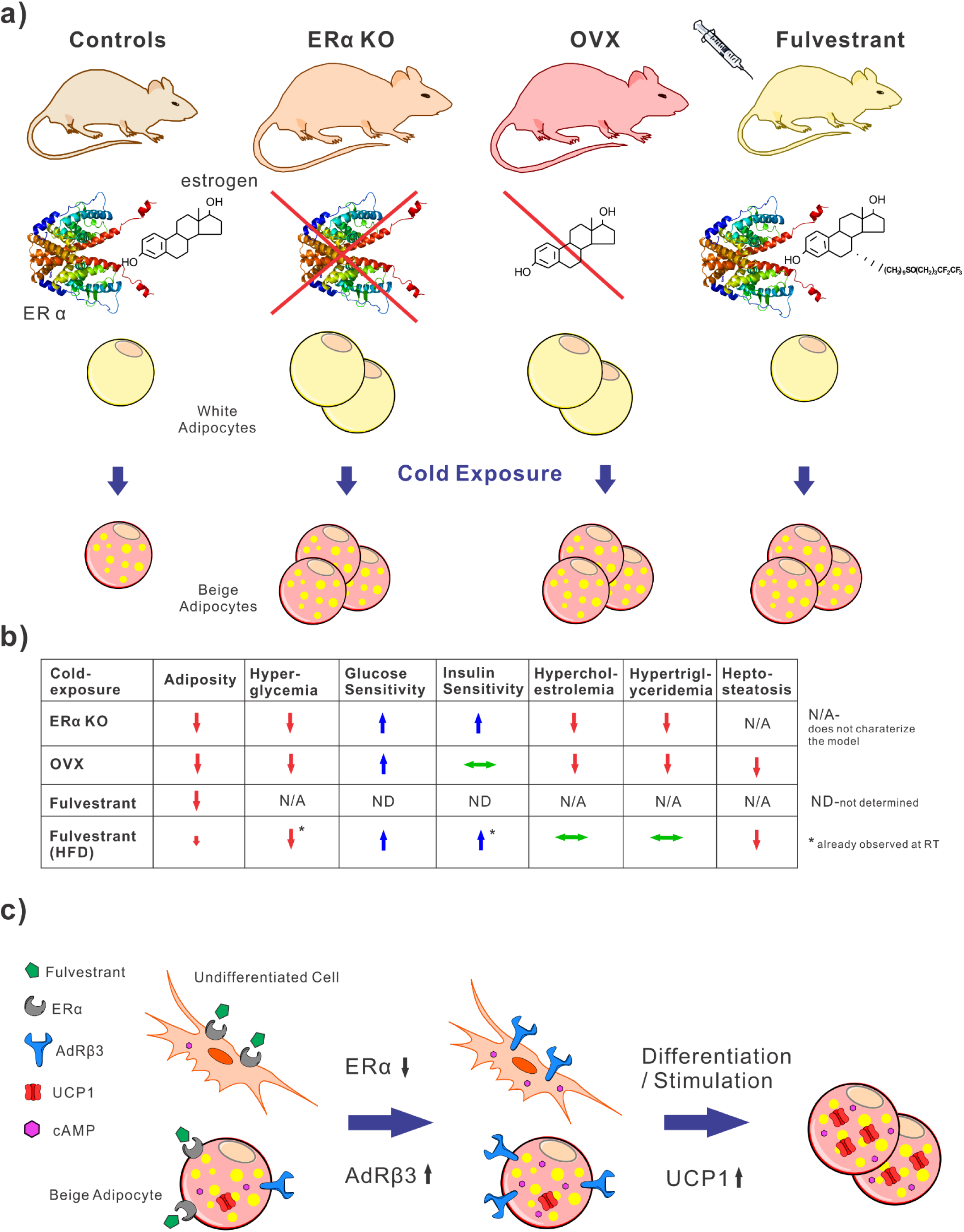
Summary. (a) In this study, we studied mouse models of inhibited ERα signaling: 1) a genetic approach - whole-body ERαKO mice; 2) a surgical approach - OVX females; 3) a pharmacological approach - treatment with the estrogen receptor antagonist – Fulvestrant, and exposed them to cold. Upon cold-exposure, we detected adipocyte beiging in control mice (e.g. WT, Sham or vehicle-treated) as expected, however, mouse models of inhibited ERα showed an enhanced response. This response included increased emergence of multilocular beige cells and elevated gene expression of beige/brown adipocyte markers in WAT and BAT. By using a primary beige cell culture, we demonstrated that this enhanced beiging is cell-autonomous. (**b**) Although under normal circumstances of room temperature, ERαKO, OVX and mice on HFD are obese and diabetic, cold-exposure resulted in significantly improved metabolism. As a result of cold-exposure, we observed: reduced adiposity and hyperglycemia, increased glucose and insulin sensitivities, and corrected diabetes-associated hypercholesterolemia, hypertriglyceridemia and hepatosteatosis. (**c**) Hypothesis – estrogen receptor inhibition enhances adipocyte beiging by “sensitizing” the cells to the beiging process upon stimulation. Stimuli of beiging may be elicited by cold-exposure or adrenergic signals. Our preliminary results support such a hypothesis – Fulvestrant pre-treatment increased cAMP levels and AdRβ3 expression in both immature and mature cells. This allowed increased beige cell differentiation or activation via AdRβ3. The mechanism that underlies enhanced cold-induced adipocyte beiging following ERα inhibition is yet to be elucidated.

## Acknowledgements

This study was supported by NIH, NIDDK (R01 DK066556 and R01 DK088220) and an AHA postdoctoral fellowship grant (K.L., 16POST27250024). We thank Brianna Findley and Xiaoli Lin from the Radiology & Advanced Imaging Research Center for their assistance in conducting the glucose uptake assay.

## Author contributions

J.M.G. and K.L. conceived, designed and interpreted the experiments. K.L. performed the experiments. A.L initiated this study. E.D.B oversaw the glucose uptake assay. J.M.G. and K.L. analyzed the experiments, discussed the results, and wrote the manuscript.

## Competing financial interests

J.M.G. is a cofounder and shareholder of Reata Pharmaceuticals. The other authors declare no competing financial interests.

---

**Figure S1.**
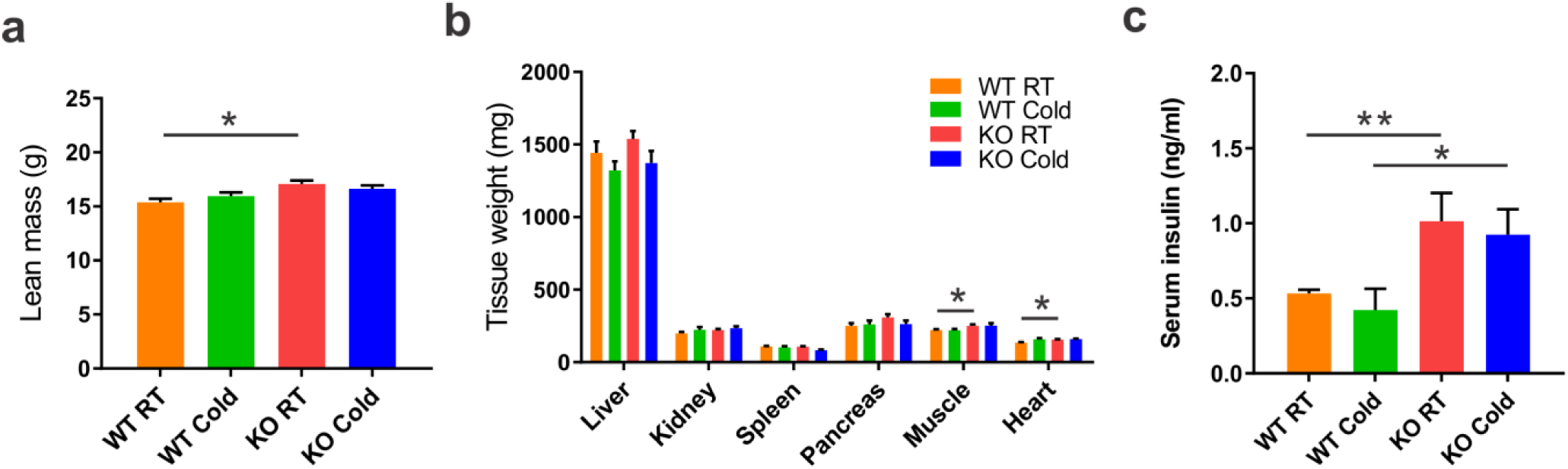
Related to Figure 1. Four to five-month-old female ERαWT (WT) and ERαKO (KO) mice were subjected to cold-exposure (6°C) for 7 days or maintained at RT (23°C). (**a**) Lean mass by NMR, n≥8. (**b**) Weight of indicated organs in WT and KO females, n≥7. (**c**) Serum insulin levels of WT and KO females, n≥5. Error bars indicate S.E.M. Statistical significance assessed by two-tailed student’s t-test, * p < 0.05, ** p <0.01.

**Figure S2.**
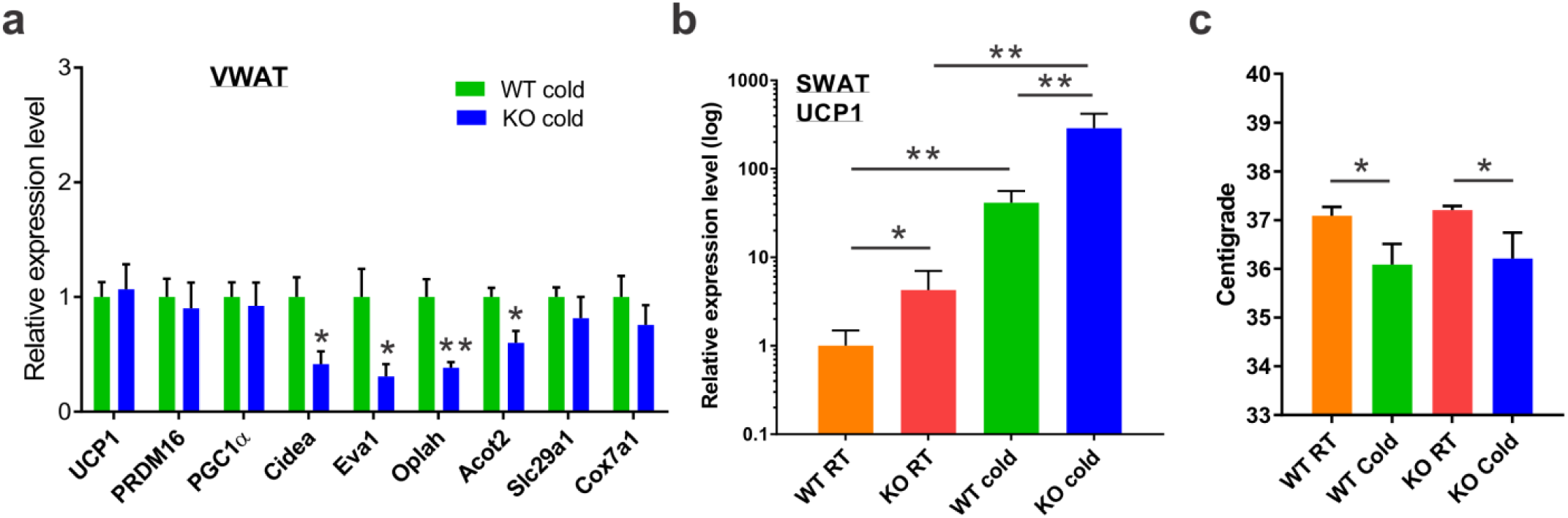
Related to Figure 2. Four to five-month-old female and male ERαWT (WT) and ERαKO (KO) mice were subjected to cold-exposure (6°C) for 7 days or maintained at RT (23°C). (**a**) Relative mRNA levels, quantified by qPCR, of brown/beige adipocyte markers expressed in VWAT (PGW) of WT and KO females at cold, n≥6. (**b**) Relative mRNA levels, quantified by qPCR, of UCP1 in SWAT (IGW) of WT and KO females at RT and cold, n≥5. (**c**) Body temperature (rectal probe) of WT and KO females at RT and cold, n≥9. Scale bars = 100 μm. Error bars indicate S.E.M. Statistical significance assessed by two-tailed student’s t-test, * p < 0.05, ** p <0.01.

**Figure S3.**
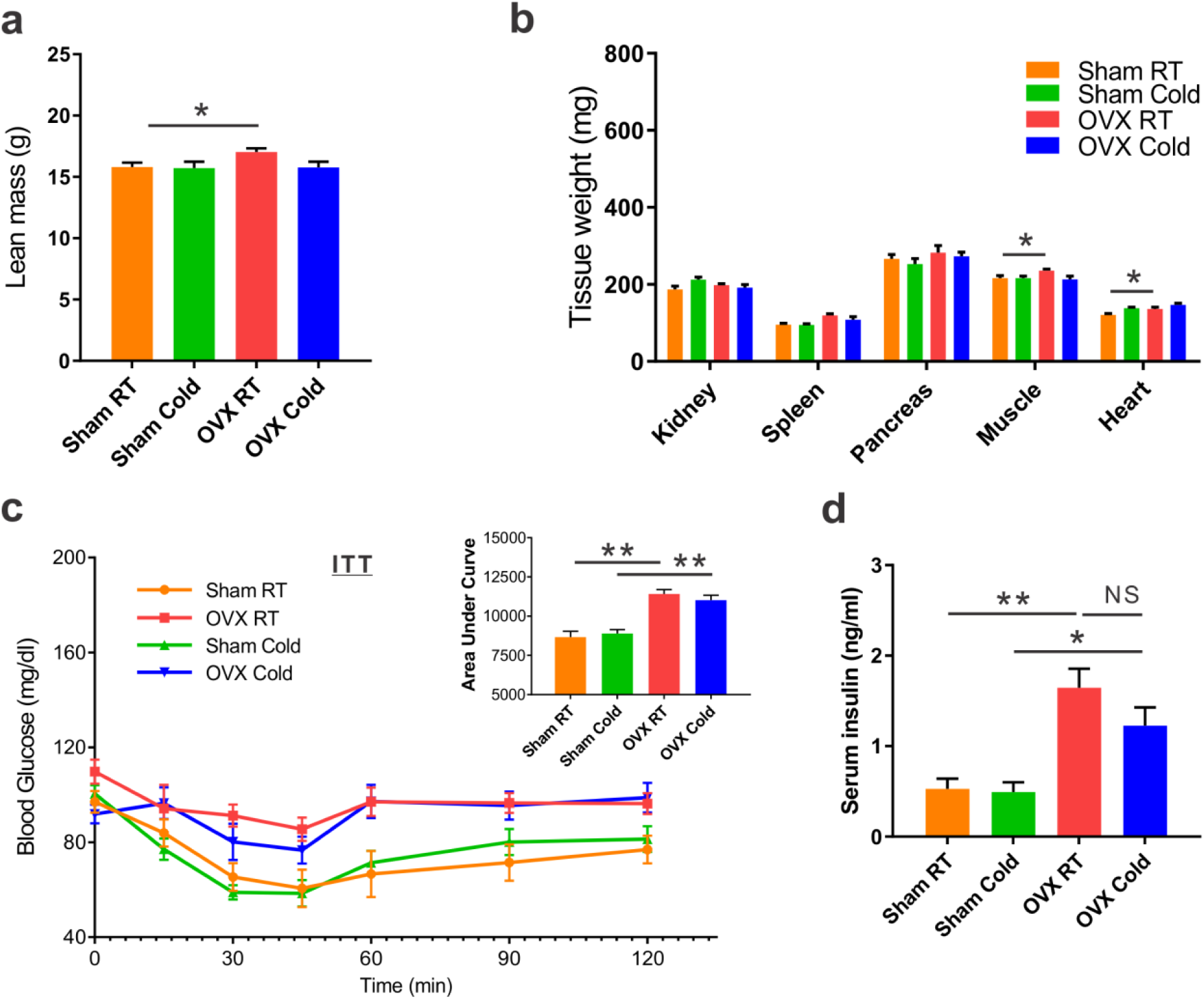
Related to Figure 3. Two-month-old female mice underwent sham operation or ovariectomy (OVX), 3 months later both groups were either subjected to cold-exposure (6°C) for 7 days or maintained at RT (23°C). (**a**) Lean mass by NMR, n≥6. (**b**) Weight of indicated organs in Sham and OVX females, n≥8. (**c**) Insulin tolerance tests were performed in 6-month-old Sham and OVX females at RT (a week prior to cold-exposure) and immediately after cold-exposure. Mice were fasted, i.p. injected with 0.75U/kg insulin, and their glucose levels were monitored, n≥10. Inset – areas under curve. (**d**) Serum insulin levels of Sham and OVX females, n≥5. Error bars indicate S.E.M. Statistical significance assessed by two-tailed student’s t-test, * p < 0.05, ** p <0.01.

**Figure S4.**
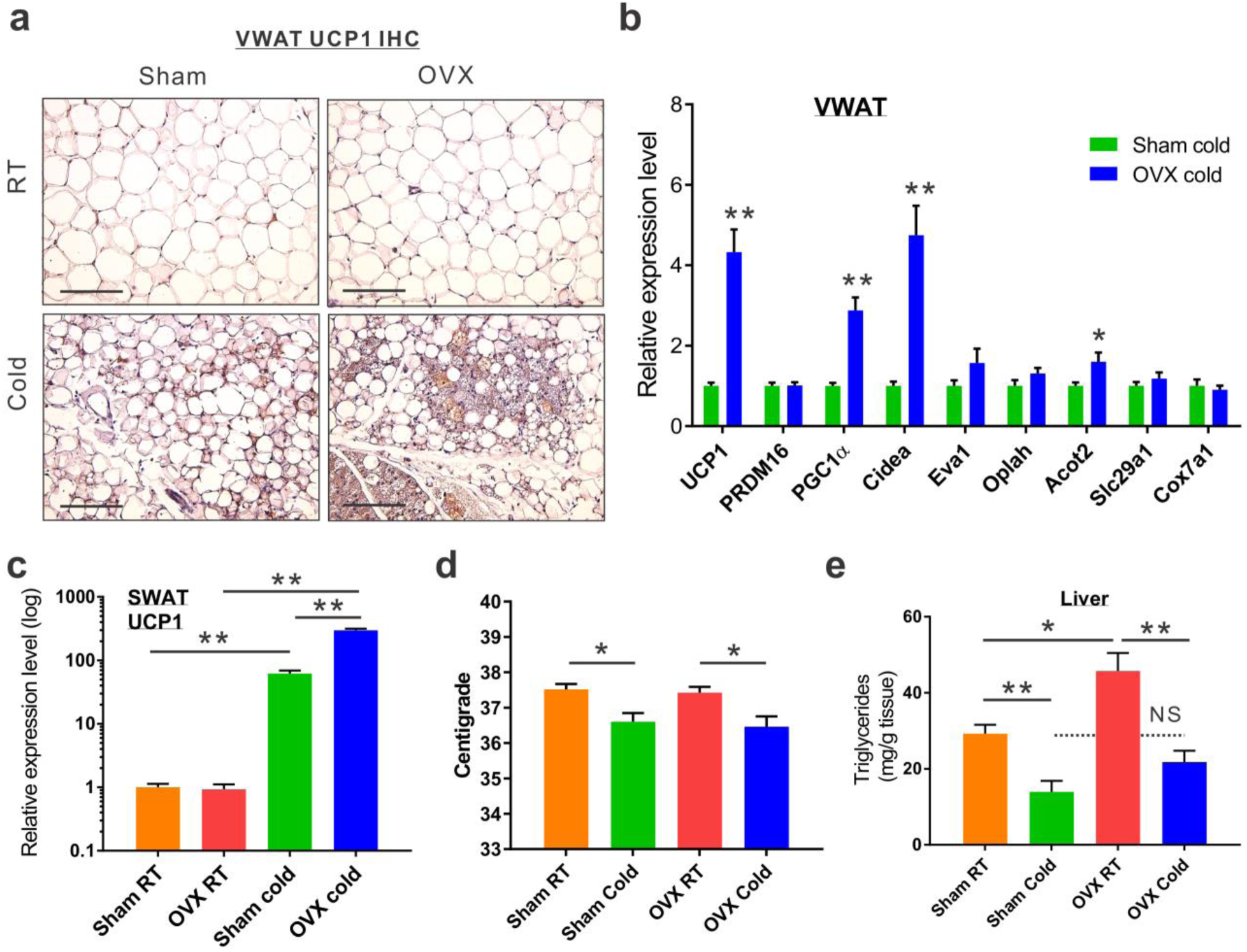
Related to Figure 4. Two-month-old female mice underwent sham operation or ovariectomy (OVX), 3 months later both groups were either subjected to cold-exposure (6°C) for 7 days or maintained at RT (23°C). (**a**) UCP1 immunohistochemistry - representative sections of VWAT. (**b**) Relative mRNA levels, quantified by qPCR, of brown/beige adipocyte markers expressed in VWAT (PGW) at cold, n≥10. (**c**) Relative mRNA levels, quantified by qPCR, of UCP1 in SWAT (IGW) of Sham and OVX females at RT and cold, n≥7. (**d**) Body temperature (rectal probe) of Sham and OVX females at RT and cold, n≥11. (**e**) Triglyceride levels in the livers of Sham and OVX females at RT and cold, n≥8. Scale bars = 100 μm. Error bars indicate S.E.M. Statistical significance assessed by two-tailed student’s t-test, * p < 0.05, ** p <0.01.

**Figure S5.**
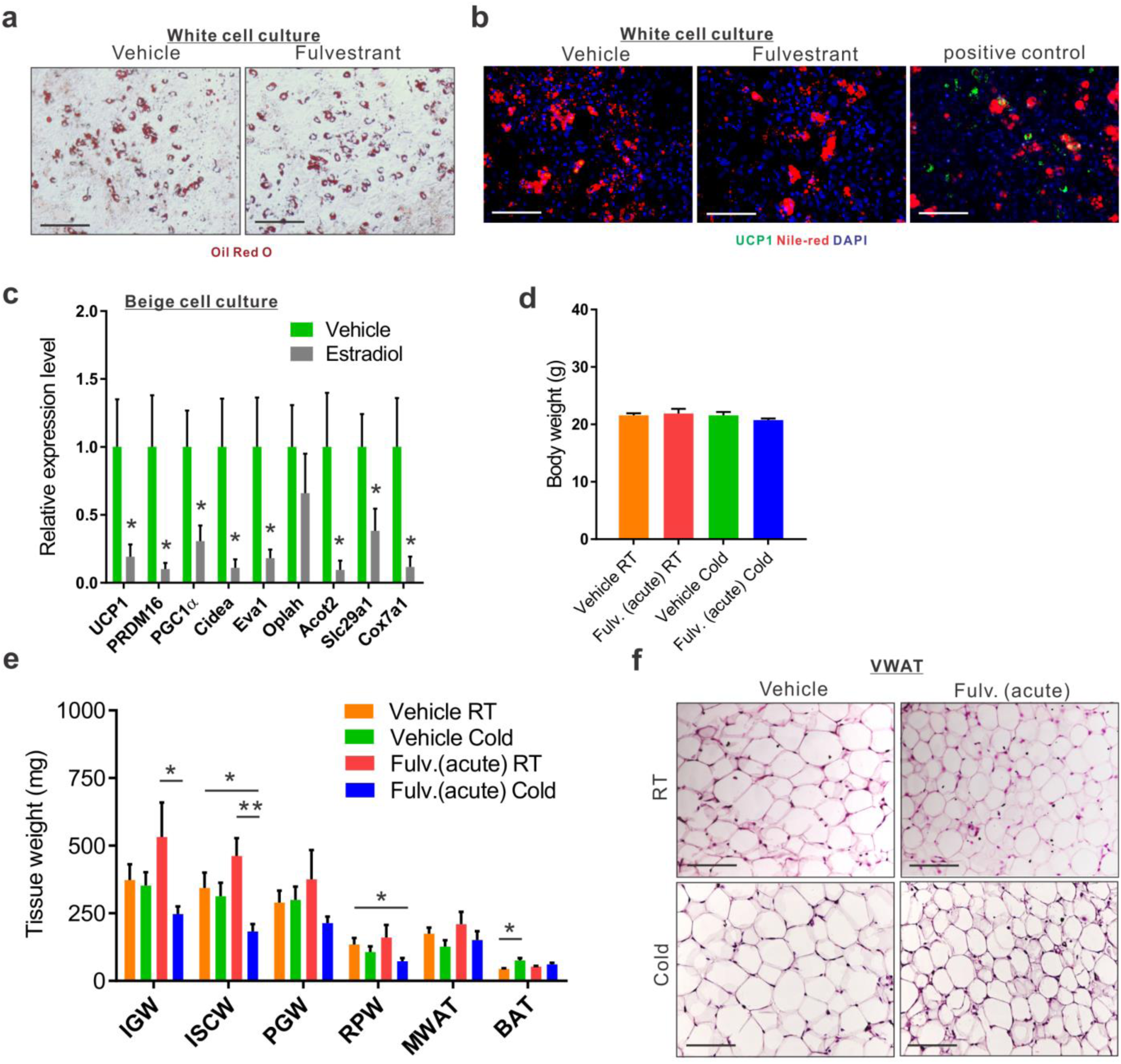
Related to Figure 5. (**a-c**) SV cells were isolated from SWAT of two-month-old females. (**a-b**) Confluent cells were induced with white adipogenic media in the presence of vehicle or Fulvestrant. A week later, cells were stained with Oil Red O (a) or immunostained for UCP1 (b). Nile Red stains lipid droplets. Positive control – confluent cells induced with beige adipogenic media for a week, and immunostained for UCP1. (**c**) Confluent cells were induced with beige adipogenic media in the presence of vehicle or Estradiol. A week later, beige cells were activated with Forskolin. Beiging was assessed by relative mRNA levels, quantified by qPCR, of brown/beige adipocyte markers, n≥6. (**d-e**) Acute administration: Four-month-old WT females were given a vehicle or 40 mg/kg/injection Fulvestrant as described in (Fig. 5d), after they were subjected to cold-exposure (6°C) for 7 days or maintained at RT (23°C). (**d**) Body weight, n≥7.(e) Weight of indicated fat depots in vehicle-treated and Fulvestrant-treated females: SWAT - IGW and ISCW; VWAT - PGW, RPW and MWAT; and intrascapular BAT, n≥7. (**f**) Representative H&E-stained histological sections of SWAT. Scale bars = 100 μm. Error bars indicate S.E.M. Statistical significance assessed by two-tailed student’s t-test, * p < 0.05, ** p <0.01.

**Figure S6.**
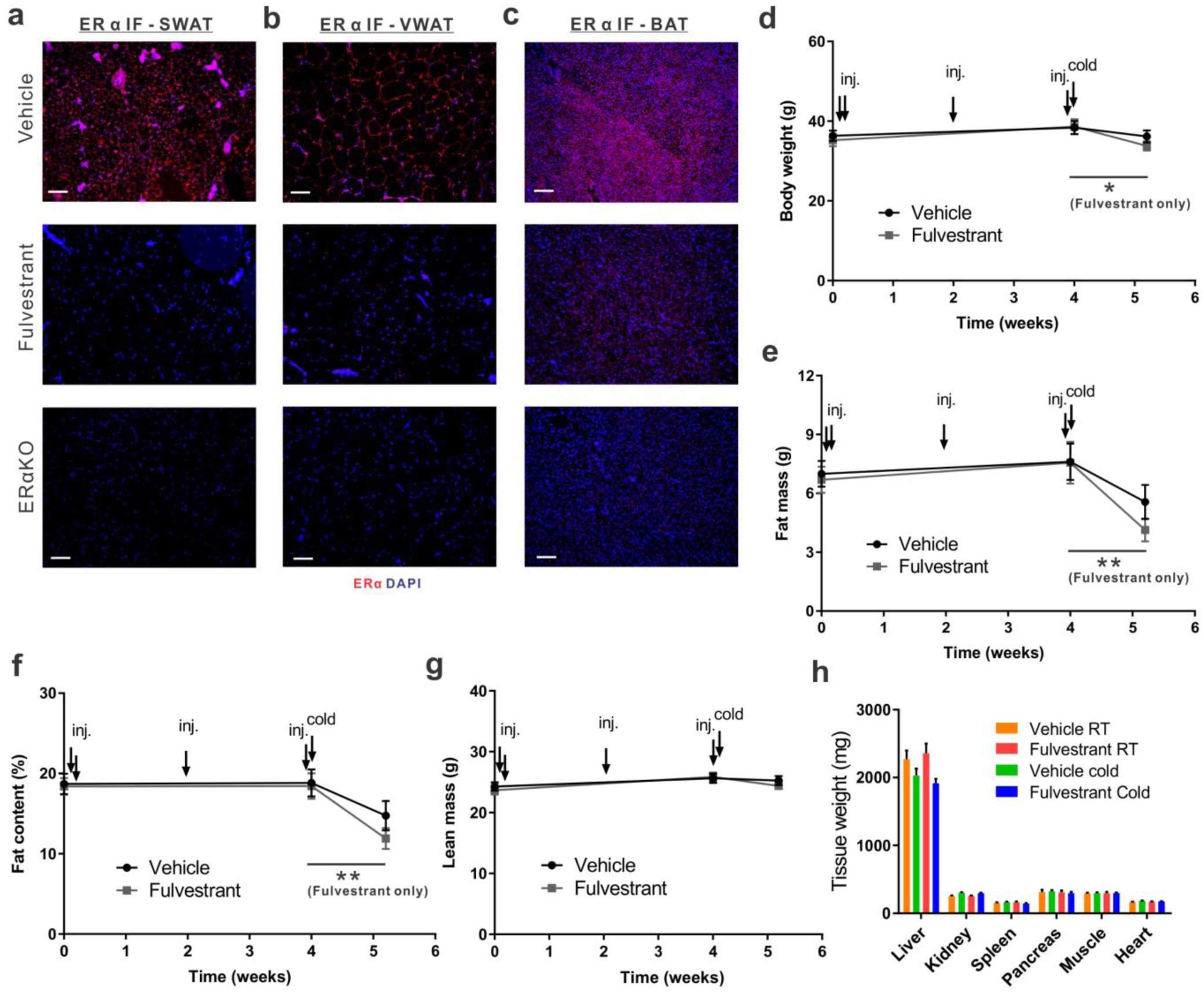
Related to Figure 6. hronic administration: Four-month-old WT ICR(CD1) females were given a vehicle or 40 mg/kg/injection Fulvestrant for a month as described in (Fig. 6a), after they were subjected to cold-exposure (6°C) for 7 days or maintained at RT (23°C). **(a-c)** ERα immunofluorescence of vehicle-treated and Fulvestrant-treated females at RT (prior to cold-exposure) - representative sections of SWAT (a), VWAT (b) and BAT (c). Negative controls (lower lane) – representative sections from ERαKO females that lack ERα expression. **(d-g)** Body weight (d), fat mass by NMR (e), fat content by NMR (f) and lean mass by NMR (g) were monitored during vehicle and Fulvestrant administration and following cold-exposure, n≥11. (**h**) Weight of indicated organs in vehicle-treated and Fulvestrant-treated females, n≥6. Scale bars = 100 μm. Error bars indicate S.E.M. Statistical significance assessed by two-tailed student’s t-test, * p < 0.05, ** p <0.01.

**Figure S7.**
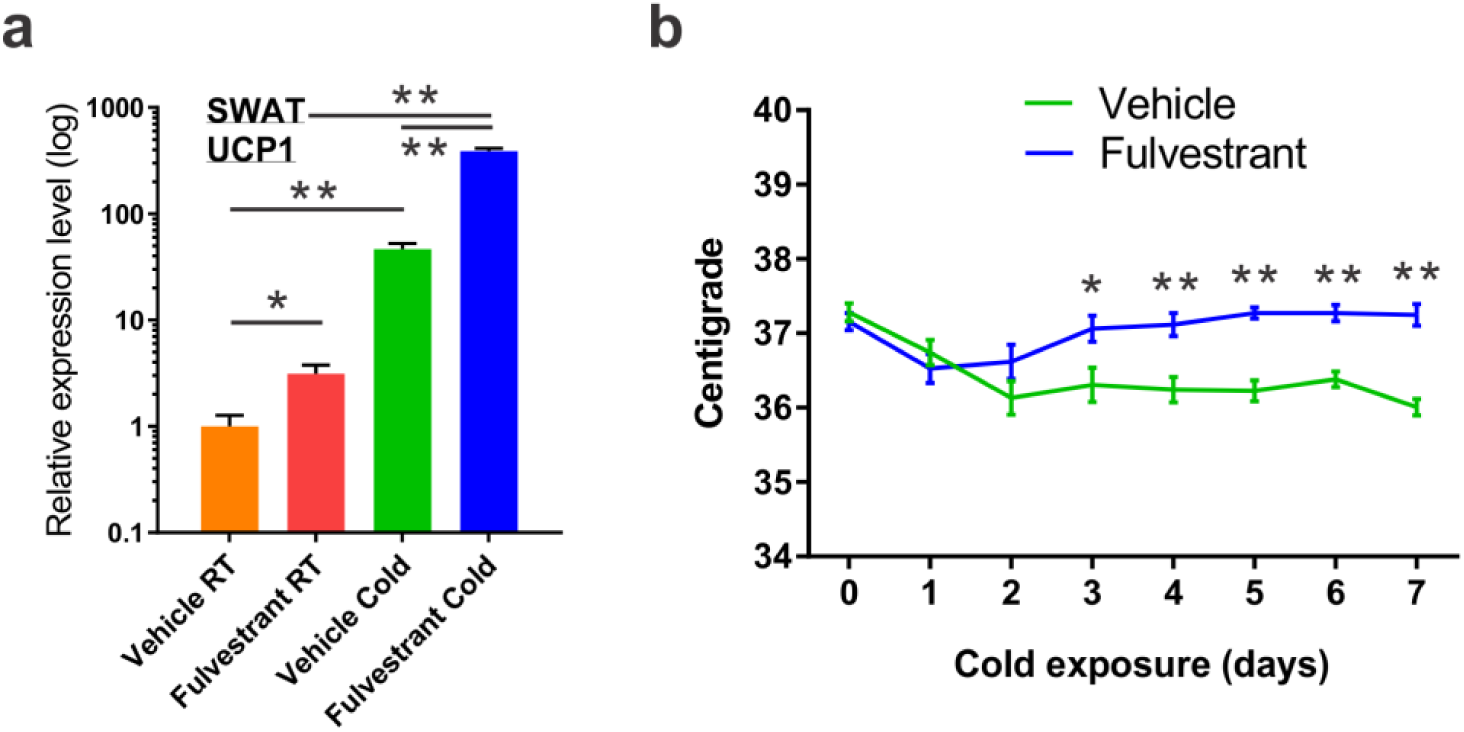
Related to Figure 7. Chronic administration: Four-month-old WT ICR(CD1) females were given a vehicle or 40 mg/kg/injection Fulvestrant for a month as described in (Fig. 6a), after they were subjected to cold-exposure (6°C) for 7 days or maintained at RT (23°C). (**a**) Relative mRNA levels, quantified by qPCR, of UCP1 in SWAT (IGW) of vehicle-treated and Fulvestrant-treated females at RT and cold, n≥6. (**d**) The body temperatures (rectal probe) of vehicle-treated and Fulvestrant-treated females were monitored upon cold-exposure, n≥12. Error bars indicate S.E.M. Statistical significance assessed by two-tailed student’s t-test, * p < 0.05, ** p <0.01.

**Figure S8.**
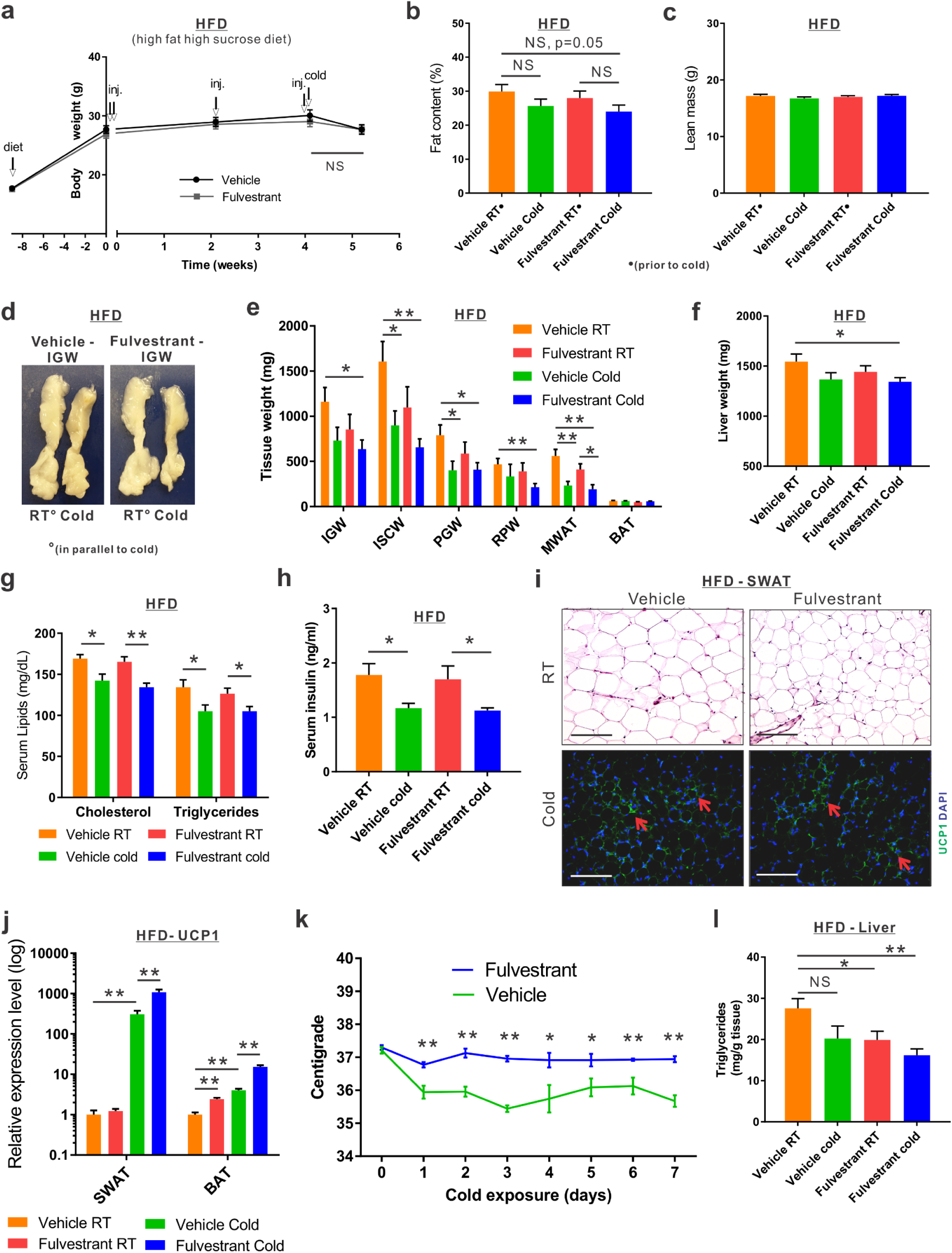
Related to Figure 8. (**a-l**) Chronic administration in a high-fat diet (HFD) model: At the age of two-month-old onwards, WT female mice were fed with high-fat high-sucrose diet. Two months later, the females were given a vehicle or 40 mg/kg/injection Fulvestrant for a month as described in (Fig. 8a), after they were subjected to cold-exposure (6°C) for 7 days or maintained at RT (23°C). (**a**) Body weight was monitored from the onset of HFD, during vehicle and Fulvestrant administration and following cold-exposure, n≥21. (**b**) Fat content by NMR, n≥17. (**c**) Lean mass by NMR, n≥17. (**d**) Representative photographs of IGW adipose depots of vehicle-treated and Fulvestrant-treated females on HFD at RT and cold. (**e**) Weight of indicated fat depots in vehicle-treated and Fulvestrant-treated females on HFD: SWAT - IGW and ISCW; VWAT - PGW, RPW and MWAT; and intrascapular BAT, n≥7. (**f**) Liver weights of vehicle-treated and Fulvestrant-treated females on HFD, n≥7. (**g**) Serum cholesterol and triglyceride levels of vehicle-treated and Fulvestrant-treated females on HFD, n≥7. (**h**) Serum insulin levels vehicle-treated and Fulvestrant-treated females on HFD, n≥7. (**i**) Representative histological sections of SWAT - H&E-stained at RT (upper lane) and UCP1 immunofluorescence at cold (lower lane). Arrows indicate UCP1+ cells. (**j**) Relative mRNA levels, quantified by qPCR, of UCP1 in SWAT (IGW) and BAT of vehicle-treated and Fulvestrant-treated females on HFD at RT and cold, n≥7. (**k**) The body temperatures (rectal probe) of vehicle-treated and Fulvestrant-treated females on HFD were monitored upon cold-exposure, n≥7. (**l**) Triglyceride levels in the livers of vehicle-treated and Fulvestrant-treated females on HFD at RT and cold, n≥7. Scale bars = 100 μm. Error bars indicate S.E.M. Statistical significance assessed by two-tailed student’s t-test, * p < 0.05, ** p <0.01.

**Figure S9.**
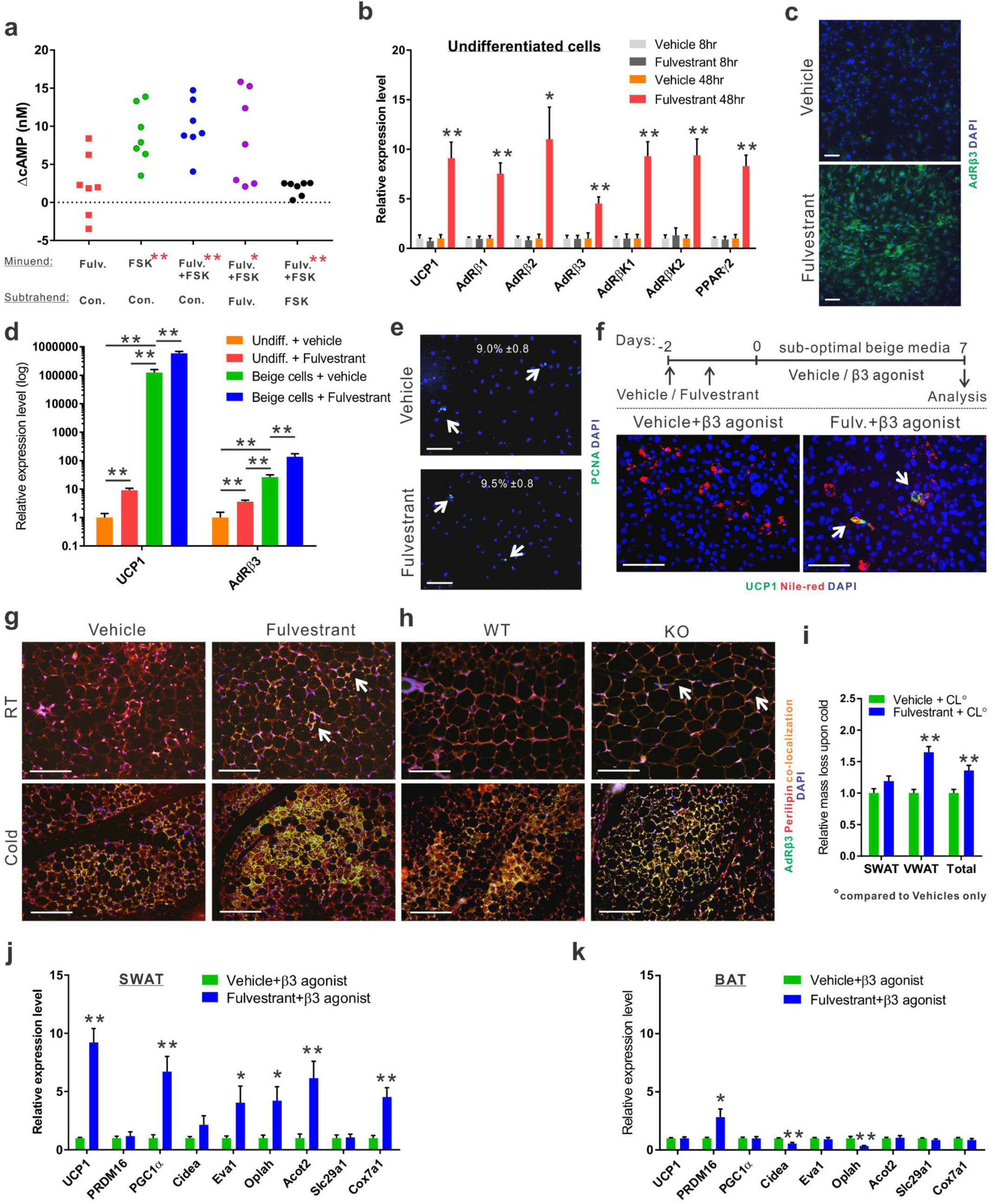
Related to Figure 9. (**a-e**) SV cells were isolated from SWAT of two-month-old WT females. Undifferentiated cells: confluent cells were treated daily with a vehicle or Fulvestrant for 8-48hr, in the absence of beige adipogenic reagents (a-e). Beige cells: Confluent cells were induced with beige adipogenic media in the presence of vehicle or Fulvestrant for a week (c). (**a**) Relative changes in cAMP levels upon induction with or without Forskolin. Every dot represents an average of technical triplicates of one biological sample. The relative changes in cAMP levels are calculated as differences between treatments, n≥7. (**b, d**) Relative mRNA levels, quantified by qPCR, of UCP1, β-adrenergic receptors, β-adrenergic receptor kinases and PPARγ2, n≥5. (**c, e**) Vehicle-treated or Fulvestrant-treated undifferentiated cells were immunostained for AdRβ3 (c) or PCNA (e) expression. Arrows indicate PCNA^+^ cells and the numbers represent their percentage.(f) Confluent cells were treated daily with a vehicle or Fulvestrant for 48hr, and then induced with beige adipogenic media in the presence of β3 agonist without Fulvestrant. Beiging was assessed by UCP1 immunostaining. Nile Red stains lipid droplets. (**g-h**) Vehicle-treated and Fulvestrant-treated females (g, see Fig. 5) as well as ERαWT (WT) and ERαKO (KO) females (h, see Fig. 1-2) were subjected to cold-exposure (6°C) for 7 days or maintained at RT (23°C). Representative histological sections of SWAT, which were immunostained for AdRβ3 and Perilipin. Arrows indicate AdRβ3+ adipocytes in Fulvestrant-treated females at RT, KO females at RT, and all specimens at cold. (**i-k**) Four-month-old WT females were pre-treated with a vehicle or 40 mg/kg/injection Fulvestrant, followed by administration of 1 mg/kg/day β3 agonist as described in (Fig. 9g). (**i**) Relative fat mass loss in SWAT and VWAT compartments were calculated according to white adipose depot weights (not shown), n≥8. (**j-k**) Relative mRNA levels, quantified by qPCR, of brown/beige adipocyte markers expressed in SWAT (IGW) (j), and BAT (k), n≥7. Scale bars = 100 μm. Error bars indicate S.E.M. Statistical significance assessed by twotailed student’s t-test, * p < 0.05, ** p <0.01. In (a), results are based on a matched standard curve and a linear regression analysis.

**Supplementary Table 1.**
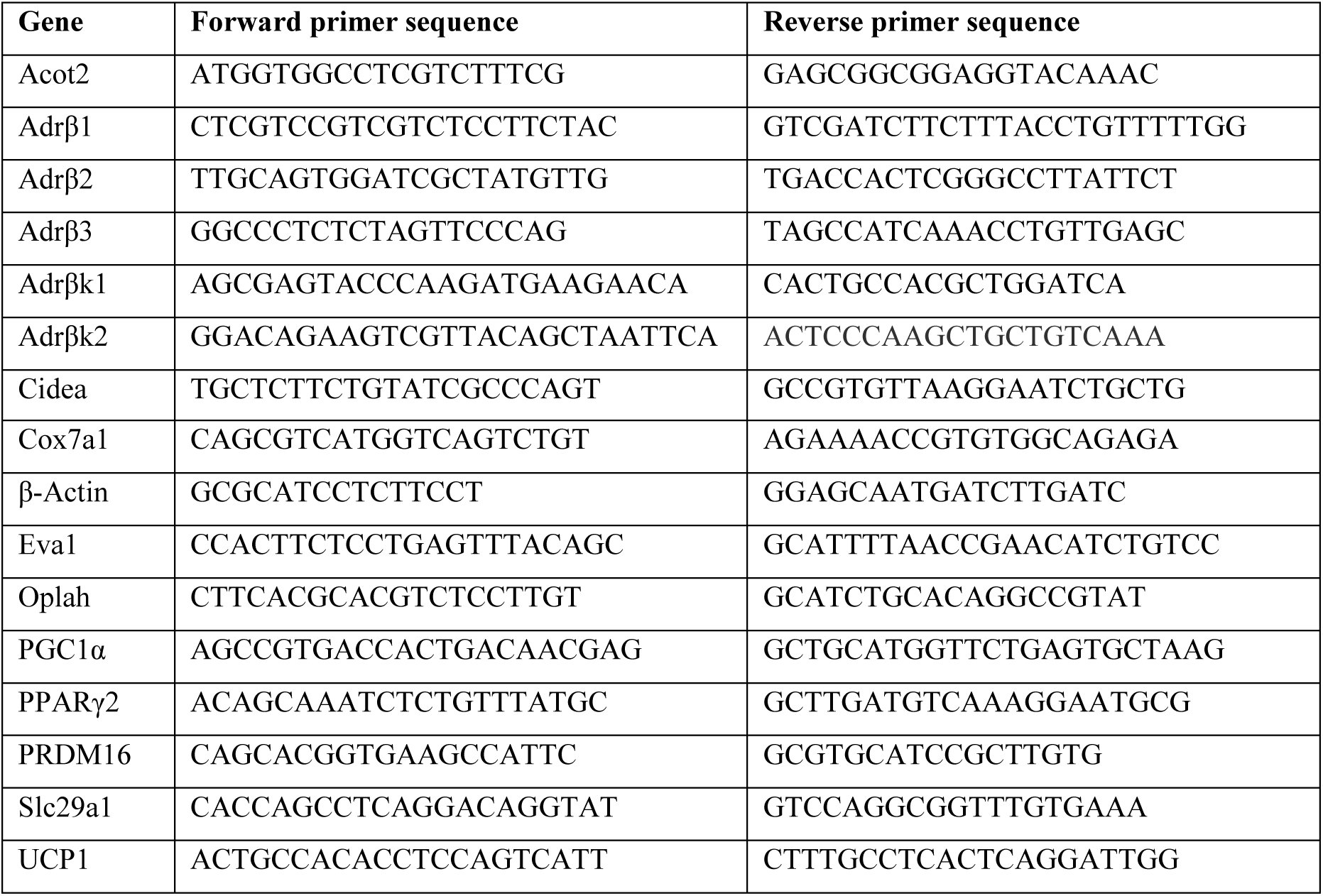
qPCR primer sequences.

